# Proteomic landscape of tunneling nanotubes reveals CD9 and CD81 tetraspanins as key regulators

**DOI:** 10.1101/2022.12.21.521537

**Authors:** Roberto Notario Manzano, Thibault Chaze, Eric Rubinstein, Esthel Penard, Mariette Matondo, Chiara Zurzolo, Christel Brou

## Abstract

Tunneling nanotubes (TNTs) are open actin- and membrane-based channels, connecting remote cells and allowing direct transfer of cellular material (e.g. vesicles, mRNAs, protein aggregates) from cytoplasm to cytoplasm. Although they are important especially in pathological conditions (e.g., cancers, neurodegenerative diseases), their precise composition and their regulation were still poorly described. Here, using a biochemical approach allowing to separate TNTs from cell bodies and from extracellular vesicles and particles (EVPs), we obtained the full composition of TNTs compared to EVPs. We then focused to two major components of our proteomic data, the CD9 and CD81 tetraspanins, and further investigated their specific roles in TNT formation and function. We show that these two tetraspanins have distinct non-redundant functions: CD9 participates in stabilizing TNTs, whereas CD81 expression is required to allow the functional transfer of vesicle in the newly formed TNTs, possibly by regulating docking to or fusion with the opposing cell.

## Introduction

Tunneling nanotubes (TNTs) are thin membranous conduits, supported by F-actin that form continuous cytoplasmic bridges between cells over distances ranging from several hundred nm up to 100 µm [1,2]. They allow cell-to-cell communication by facilitating the transfer of different cargoes directly from cytoplasm to cytoplasm of the connected cells, including organelles (e.g., lysosomes, mitochondria) [3–5], micro- or mRNAs, pathogens [6,7] and misfolded/aggregated proteins (e.g., prion proteins, tau or α-synuclein aggregates [8–10]). TNTs could play major roles in various diseases, including neurodegenerative diseases [11–13] or cancers of different types [14]. In addition to cell cultures and tumors explants [14], TNT-like connections have been shown to exist in the retina and facilitate cellular material transfer between photoreceptors [15,16] or pericytes [17], highlighting the importance of understanding the biology of these protrusions in order to unravel their possible role(s) *in vivo* [12]. TNT formation is highly dynamic and appears to be regulated by cellular stresses and actin regulators [18]. Two models have been proposed for TNT formation. In the first, TNTs would be produced by membrane deformation and elongation of the protrusion supported by actin polymerization followed by adhesion and fusion of this protrusion with the opposing cell. Alternatively, TNTs could be formed by cell dislodgement where adhesion and membrane fusion would be the first steps before cell bodies separation and elongation of the protrusion by actin polymerization [18,19]. However, TNTs have been shown to be structurally more complex, being made up of bundles of fine open connections (called iTNTs) held together by partially identified proteins, including N-Cadherin [2]. The mechanism(s) and specific pathways governing iTNT/TNT formation as well as the molecular components of TNTs are still not known.

In addition to TNTs, one of the major pathways used by cells to transfer materials over long distances is through membrane-surrounded vesicles, collectively known as extracellular vesicles (EVs) [20]. EVs are released by all cells, and up taken by distant recipient cells [21–23]. They can be formed either by direct budding from the plasma membrane, or by secretion of intraluminal vesicles of multivesicular compartments (in which case they are called exosomes). Because of their common functions, their similar diameters, and because TNTs are fragile and easily broken and therefore can be released in cell culture supernatants (where EVs are also found), it has been challenging to distinguish TNTs from EVs in terms of composition [24]. In this regard, two members of the tetraspanin family, CD9 and CD81, which are well-known and widely used markers for EVs [25], have been detected in TNTs in T cells when overexpressed [26].

Tetraspanin form a family (with 33 members in mammals) of small four membrane-spanning domain proteins with two extracellular domains, including a large one harboring a tetraspanin-specific folding and two short cytoplasmic tails. Tetraspanins are involved in various cellular processes like migration, adhesion, signaling, pathogen infection, membrane damage reparation, membrane protrusive activity and cell-cell fusion [27–30]. Their function is linked at least in part to their ability to interact with other transmembrane proteins, forming a dynamic network or molecular interactions referred to as the tetraspanin web or Tetraspanin-enriched microdomains (TEM) [31]. Inside this “web”, the tetraspanins CD9 and CD81 directly interact with the Ig domain proteins CD9P1 (aka EWI-F, encoded by the PTGFRN gene) and EWI-2 (encoded by the IGSF8 gene) [32–35], which have an impact on several fusion processes [36–38]. Whether CD9 and CD81 are also endogenously present in TNTs from non-T lymphoid cells, and whether they play a role in the formation or function of TNTs has not been investigated.

With the goal of identifying structural components of TNTs, and possibly specific markers and regulators of these structures, we established a protocol of TNT isolation from U2OS cultured cells [39], allowing to separate TNTs from extracellular vesicles and particles (EVPs) and from cell bodies. We obtained the full protein composition of TNTs, compared to EVPs. As CD9 and CD81 were major components of TNTs, we further studied their specific roles in TNT formation and ability to transfer cellular material using human neuronal SH-SY5Y, a well-known cell model to study functional TNTs [2,40]. Our data indicate that CD9 and CD81 have different and complementary functions by possibly regulating two different steps of the formation of TNTs.

## Results

### Purification of TNTs and EVPs

In order to isolate nanotubes and similar structures (generically referred to hereafter as TNTs), we took advantage of the fact that TNTs are very sensitive to mechanical stress as they are not attached to substrate [1,41]. We used U2OS cells because they are robustly adherent cells exhibiting few long protrusions and are able to grow TNTs in complete and serum-free medium and transfer cellular material through these bridges (Fig S1A, B and [42]). Cells were first plated and cultured in serum-free medium to facilitate the purification steps; next after removal of the cell medium and replacement by a small volume of PBS, the flasks were vigorously shaken to break the TNTs that were then isolated from the supernatant by ultracentrifugation after elimination of floating cells by low-speed centrifugations and filtration (see workflow on Fig 1A). When necessary, extracellular vesicles and particles (EVPs) were directly collected from the first cell culture supernatant and enriched following a standardized procedure ([25,43–45] and see Fig 1A). Observation of the obtained particles using transmission electronic microscopy (TEM) revealed that they were morphologically different (Fig 1B). EVPs appeared mostly circular with a mean diameter of 60 nm (Fig 1C), whereas TNTs (or TNT fragments) were mostly cylindrical, with mean diameter of 69 nm and length varying from 140 to 900 nm (mean length 372 nm). These results were in accordance with the expected sizes of EVPs, and with the different nature of the materials of each fraction. To further validate that our protocol for preparing TNTs vs. EVPs was accurate, we sought to engineer cells with fluorescent TNT markers. As shown in Fig S1A, actin chromobody-GFP (an actin-detecting probe that does not affect actin dynamics [46]) decorated TNTs after stable expression. Endogenous CD9 was present on at least a fraction of TNTs formed between U2OS cells (cultured in serum-free or -rich conditions, Fig S1A), and a GFP-CD9 construct also decorated these structures when stably expressed in these cells (Fig S1B). As a control, cells stably expressing H2B-GFP (labeling nuclei) were used.

**Figure 1:**
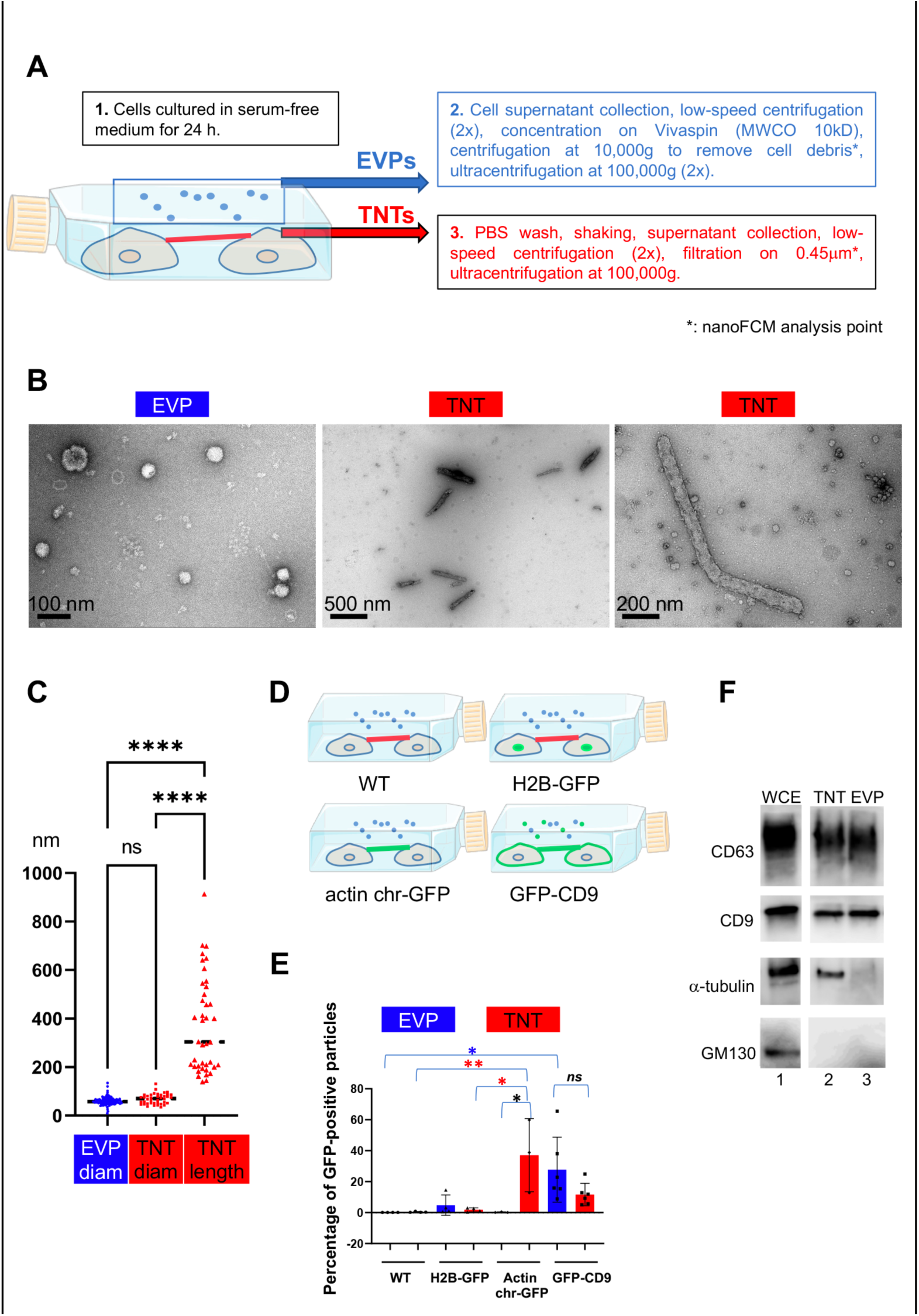
Validation of the purification procedures A. Workflow for TNT vs. EVP purification. EVPs were purified from cell culture supernatant, TNTs from remaining attached cell supernatant after shaking. * indicates the fraction used for nanoFCM analysis. B. Representative pictures of negative staining and Transmission Electron Microscopy from EVP and TNT fractions as indicated. Scale bars are 100, 500 and 200 nm respectively. C. Violin plot of the size distribution of EVP diameters (124 vesicles) and TNT diameters and lengths (40 objects), line is the median. EVP and TNT diameter means are 60 and 69 nm respectively, TNT lengths extend from 140 to 912 nm, mean at 372 nm. Statistical analysis is one-way Anova with Tukey post-hoc correction. Ns, non-significant, ****p<0.0001. D. Schematic representation of stable cell lines where green color indicates the location of GFP-tagged protein: H2B-GFP (nuclear), actin chromobody-GFP (actin cytoskeleton including TNTs) and GFP-CD9 (cell surface, TNTs, EVPs). E. Scatter dot plot representing the mean percentage (with SD) of GFP-positive particles analyzed by nanoFCM. Statistical analysis of 3 independent experiments to 6 for GFP-CD9 (oneway Anova with Tukey post-hoc correction) show the following respective p-values (from top): blue*: 0.0252, red**: 0.0089, red*: 0.0131, black*: 0.0161. Means values are (from left to right): 0.01, 0.3, 4.8, 1.75, 0.16, 37.07, 27.7 and 11.6. F. Western blot of WCE (20μg, corresponding to around 0.1×10^6^ cells), TNT and EVP fractions (both from 10 10^6^ cells) prepared from the same cells, blotted with CD63, CD9, α-tubulin and GM130 specific antibodies. White lane indicates that intervening lanes of the same gel (and same exposure) have been spliced out.

Four independent preparations of TNTs and EVPs from the same cell cultures (Fig 1D), were analyzed. We confirmed that the crude preparations (before ultracentrifugation, Fig 1A) contained particles and measured their size and their fluorescence using Nano flow-cytometry. As shown in Fig 1E and S1C-E, both EVPs and TNTs contained particles of similar mean diameter (around 60 nm, Fig S1C, D), ranging from 40 to more than 100 nm, in perfect accordance with TEM results. The sizes and concentrations of EVPs and TNTs were the same in parental and transfected cell lines (Fig S1D, E). Both fractions contained particles bearing GFP-CD9 (Fig 1E), confirming the presence of membrane-enclosed particles. A major difference between the two fractions was the large proportion of particles in the TNT preparation (38%) that bore the actin chromobody-GFP which was barely detected in EVPs (0.2%). Finally, H2B-GFP was present in a very minor part of the particles, and the Golgi marker GM130 was detected neither in EVPs nor in TNTs by western-blot (Fig 1F), showing that contaminations with cell debris or nuclei were limited, thus validating our protocol. CD9 and CD63 were detected in both TNT and EVP preparations, whereas tubulin (shown to label a fraction of TNTs on Fig S1A) was only detected in TNT fraction. The classical marker of EVPs Alix was mainly detected in EVP fraction, and the ER protein calnexin was detected in neither fraction (Fig S1F). We checked that shaking the cells did not affect their shape and plate attachment (Fig S1G, 2 left pictures). In contrast, it decreased the percentage of connections between cells, identified after mild trypsinization and phase image analysis of fixed cells (right pictures and graph of Fig S1G) cultured in serum-free medium. To verify that culture of the cells in serum-free medium had no consequence on the nature of the TNTs, we compared the number of connection-forming cells in both conditions (Fig S1G), as well as the ability of cells to transfer DiD-labeled vesicles to acceptor cells in a cell contact-dependent manner (Fig S1H). We observed that connections were formed in both culture conditions, and that cell-to-cell transfer was happening in a similar ratio in serum-rich and reduced conditions. Together with the expression of several markers in both conditions on these cells (Fig S1 and S2), these results confirmed that serum deprivation did not affect the nature of the TNTs. Altogether, these data supported that our protocol allowed differential enrichment of the fractions in TNTs and EVPs respectively.

### Analysis of TNTome

To obtain a full and accurate picture of U2OS TNT content, we made 12 independent preparations of TNT fractions (in red in Fig 1A), each starting from about 20×10^6^ cells, and analyzed them by LC-MS/MS. These fractions are enriched in the TNT-microsomal-type/membrane proteome, although we cannot exclude that they also contain small portions of cell bodies or ER. For simplicity we shortly call them TNTome hereafter. 1177 proteins were identified in at least 9 preparations (table S1). We first observed that proteins previously described in TNTs, like actin, Myosin10 [47], ERp29 [42], or N-cadherin [2,48] were indeed present in the TNTome. Less than 100 nuclear proteins (according to GO Cellular component analysis), i.e., less than 8%, were found, which could result from partial contamination with cellular debris or dead cells. This is in accordance with nanoFCM results, where H2B-GFP positive particles were 4% of actin-chromobody-GFP positive ones. The 1177 proteins have been ranked in 4 quartiles depending on their relative abundancy (average iBAQ), highlighting the enrichment of specific factors when considering gene ontology (table S1, Fig 2A). These factors could be structural components of TNTs, but also material that was circulating in them or in the process of being transferred at the time when TNTs were broken and purified. It could be why mitochondrial (8% of the total), lysosomal/endosomal or other vesicle proteins are listed. The Proteomap analysis (Fig 2A) also revealed that TNTome is rich in RNA-associated proteins (ribosomes, translation factors, ribonucleoproteins), in accordance with TNTs being able to transfer micro and mRNAs [49,50].

**Figure 2:**
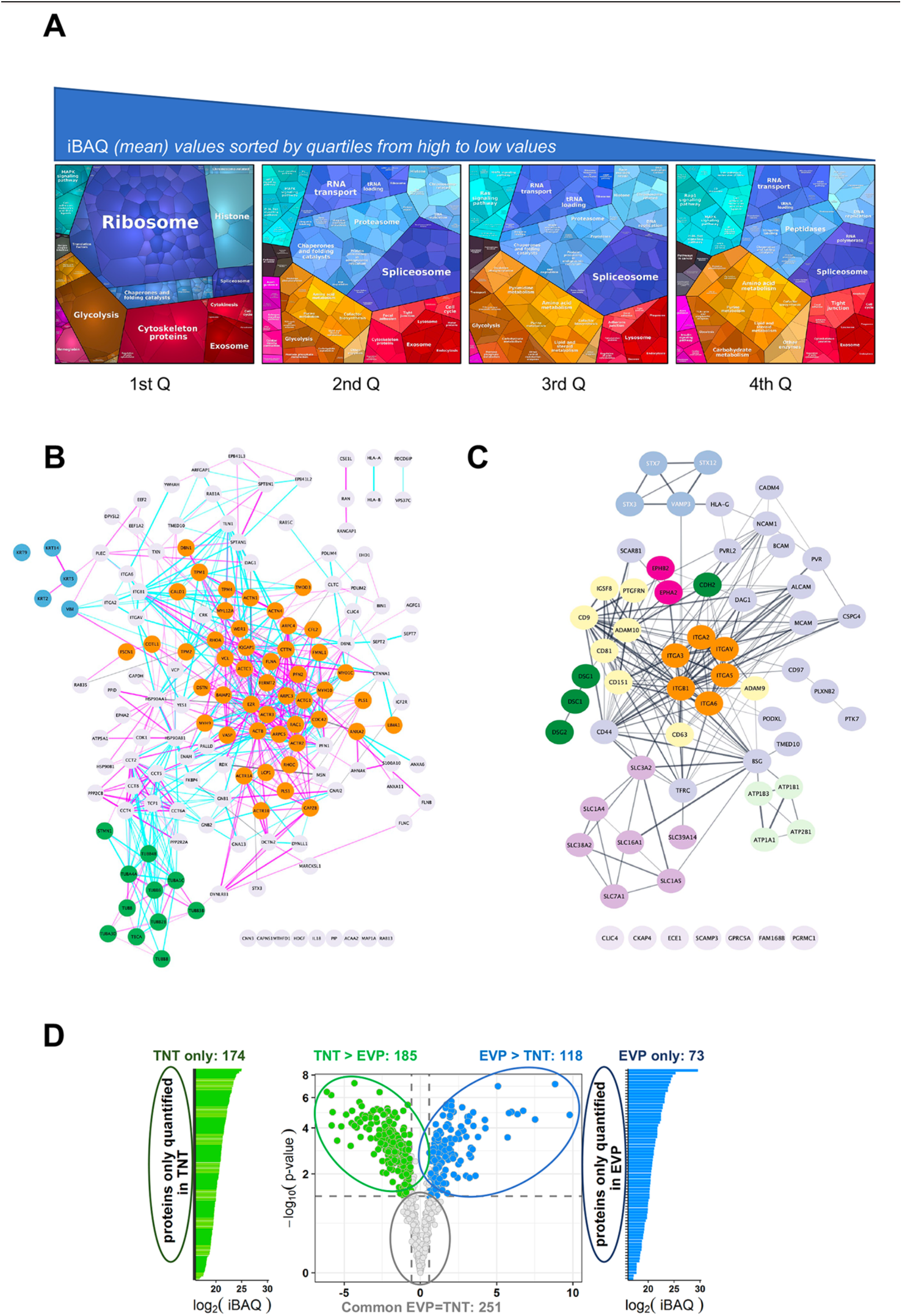
Analysis of the TNTome A. Proteomap of the 1177 proteins of the TNTome, sorted in 4 quartiles depending on their mean iBAQ. Protein accession and mean iBAQ were used to create ProteoMap, analyzed according to Gene Ontology. B. STRING physical association network for the cytoskeleton-related proteins listed in table S2. Color groups were created using Cytoscape. Green are microtubule-related proteins, blue are intermediate filaments, orange are actin-interacting proteins. Blue and pink edges show physical interactions based on databases and experiments respectively. C. Full STRING functional association network for integral surface membrane proteins of TNTome, based on table S3. Orange are Integrin proteins, red are Ephrin receptors, dark green are Cadherins, light green are Sodium/potassium transporting ATPase ions channels, purple are monocarboxylate and amino acids transporters, and yellow are tetraspanin-related proteins. D. Volcano plot of the mass spectrometry analysis based on the 4 EVP and TNT preparations, showing the maximum log2(Fold-change) in x-axis measured between TNT and EVP fractions and the corresponding -log10 (p-value) in y-axis. Dashed lines indicate differential analysis quadrants with log2 (Fold-change) =0.58 and false discovery rate FDR = 1%. Common EVP=TNT are non-significantly different (FDR >0.05) with FC>1.5, and FC<1.5. Each quadrant is named above and the number of identified proteins is indicated. Left and right are proteins non overlapping in both fractions: TNT-only and EVP only. Note that in EVP-only fraction, 10 proteins were found in TNTome (based on 12 experiments) and should therefore be removed. For the TNT proteins, only the proteins also present in TNTome have been counted.

A major group of proteins of interest were related to cytoskeleton (15%, i.e., 172 proteins, see table S2, analysis of GO terms, cellular components). As shown by STRING functional network representation, actin-related proteins were majority (Fig 2B, orange nodes) compared to microtubule-related (green nodes) or intermediate filament-related proteins (blue nodes). Actin was the 4^th^ more abundant protein of TNTome, whereas tubulin beta and alpha chains were ranked in positions 6 and 7. This was in accordance with the nature of TNTs, mainly supported by actin cytoskeleton [1]. However, recent work has shown that TNTs, in addition to actin, could contain microtubules as in some cancers (Fig S1A and [14,51]), and cytokeratins like in the case of urothelial cells [52,53].

When looking at membrane proteins, analysis of the GO terms (cellular components) of the TNTome classified around 500 proteins as membrane-related, 64 of which being strictly integral plasma membrane (see table S3 and Fig 2C). Among the latter, N-cadherin and other cadherin-related proteins (green nodes), as well as known N-cadherin interactors like alpha-catenin, were found. We also noticed the presence of various integrin subunits (orange nodes), including the Integrin β1. However, the TNTome had only a partial overlap with integrin adhesion complexes ([54], Fig S2A and tab1 of table S4) and with the consensus adhesome (Fig S2B, tab2 of table S4). In addition, very important proteins of the focal adhesions, like Paxillin, GIT2, Parvins, and PINCH1 were not in the TNTome, making unlikely focal adhesion being isolated in TNT preparations. We also analyzed whether TNT preparations could be contaminated by filopodia, another type of protrusions grown by the cells. Despite common proteins described as core filopodia proteins in U2OS cells (like Myosin10, Integrin α5, TLN1, FERMT2, MSN, LIMA1), the TNTome was devoid of others (Myo15A, TLN2, PARVA, ITGB1BP1, see [55]), ruling out an important contamination of TNTs with filopodia during the preparation. Together, these results suggested that the proteins identified in TNTome represent a specific composition of these protrusions, and not a pool of other types of protrusions which could have contaminated the preparation. We confirmed these results by immunofluorescence both for proteins found (Integrin β1, CD151, Vinculin) or not (Paxillin, GM130) in the TNTome in U2OS cells cultured in the presence and absence of serum (Fig S2C&D).

We also looked whether the TM proteins of the TNTome were the most abundant TM proteins of U2OS cells. We compared the rank in TNTome to the protein concentration [56] or to the level of RNA [57]. Whether some factors are indeed highly expressed in U2OS cells (for instance Integrin β1, SLC3A2, basigin, CLIC channels) and ranked in the first quartile of TNTome, others are weakly expressed in U2OS cells, but still enriched in TNTome (N-cadherin for instance). In addition, some cell surface proteins are highly expressed in U2OS cells (according to their corresponding RNA level [57]), but not detected in the TNTome, like SLC2A11 (Glucose transporter), APP, ITM2C, TNFR. As shown by WB in Fig S2G, we could detect membrane or membrane-associated proteins in U2OS cell extracts that were not listed in TNTome, like Integrin β4, α4, EGFR, Cx43, and APP [58]. Likewise, some proteins of very low abundancy in cells were found in TNTome (CALML5 for example), maybe reflecting a specific role in TNT formation. We also observed that the relative abundance of some proteins in TNT fractions compared to WCE (Fig S2H) was variable. As examples, CD9 and especially ADAM10 seemed to be relatively abundant in TNTs compared to WCE, whereas Integrin β1 or ANXA2 TNT/WCE ratios were much lower. Altogether, these data suggested that TNT membrane composition was not just a fraction of cell surface membrane proteins, but rather that some proteins were excluded, other more present. This first consolidated our TNT purification procedure, and second highlighted that specific mechanisms and factors should be at stake to grow and maintain TNTs.

### Comparison of the content of TNTs and EVPs

Because both EVs and TNTs have similar characteristics (membrane-formed, diameter, ability to transfer material to remote cells), we analyzed their respective composition when prepared from the same cell cultures, following the full protocol schematized in Fig 1A, from 4 independent experiments. 961 proteins in total were identified at least in 3 of the 4 TNT and EVP preparations. When keeping TNT proteins that were also present in the 1177 list of TNTome (in 9 TNT preparations over the total of 12), a total of 801 proteins were finally differentially analyzed. Our results showed a different composition of TNTs and EVPs, although common factors represent 75% of them (see volcano plot in Fig 2D). Interestingly, 174 proteins were specific for TNTs when compared to EVPs (see tab1 of table S5). Among the most abundant was the ER chaperone ERp29, previously shown to be required for the formation of TNTs [42]. When discarding organelle-associated or translation linked proteins, 89 proteins remained in this TNT-only list (table S5, tab2, constitutive). 20% of them were involved in cytoskeleton, including the positive regulator of TNTs Myosin10 [47]. When analyzing the proteins differentially abundant in TNT vs. EVPs (Fig 2D and table S6), we noticed the enrichment of cytoskeleton-related proteins, especially actin, in TNTs compared to EVPs.

The tetraspanins CD9, CD81 and CD63, classical markers of EVs were also detected in TNTs, CD9 and CD81 being among the most abundant transmembrane proteins of TNTome (table S3). While CD9 and CD63 were more abundant in EVs than in TNT, CD81 was present at similar level in both preparations. Their direct interacting partners CD9P1 and EWI2 (respectively PTGFRN and IGSF8 in Fig 2C) were present in TNTome, but more enriched in EVPs than TNTs. Consistent with the presence of the integrins α3β1 and α6β1, the tetraspanin CD151 which directly associates with these integrins was also detected in TNTome [59]. Finally, the presence of CD9, CD81, and CD151 in TNTs was confirmed by immunofluorescence in U2OS (Figs S1A and S2C).

### Differential regulation of TNT number by CD9 and CD81

To study the role of CD9 and CD81 in the formation of TNTs, we decided to use a cellular model in which TNT structure and function have already been largely investigated, SH-SY5Y cells. These human neuronal cells form many TNTs in complete medium (about 30 % of WT cells are connected by TNTs), which, contrary to U2OS cell TNTs, can be easily quantified, and distinguished from other protrusions by fluorescent imaging [2,10,60]. Similar to what we observed in U2OS cells, and in accordance with the composition of the TNTome, CD9, CD81, CD151, Integrin β1 and vinculin were detected on protrusions connecting SH-SY5Y cells (Fig S2C & E, Fig 3A), whereas TNTs appeared mostly free of Paxillin, and of the Golgi marker GM130 (Fig S2E), as it was the case for U2OS cells. Importantly, all protrusions connecting two cells, containing actin and not attached to the substrate, which correspond to the TNT morphological definition [1], bore both CD9 and CD81, indicating that these tetraspanins are good markers for TNTs (Fig 3A).

**Figure 3:**
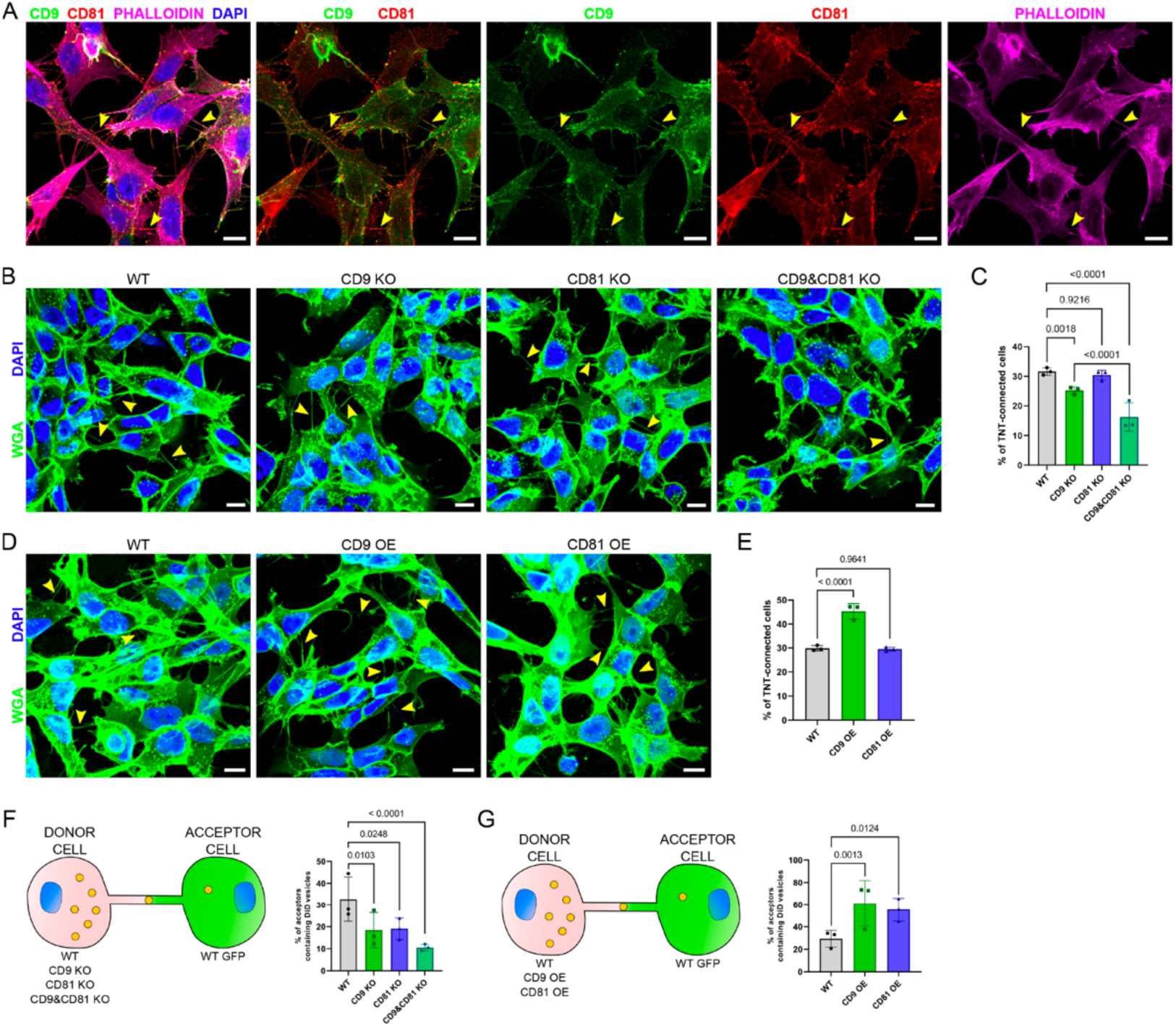
Expression of CD9/CD81 in TNTs and effects of their overexpression or invalidation A. Immunofluorescence of CD9 (green) and CD81 (red) in SH-SY5Y cells. Cells were also stained with phalloidin (magenta) and DAPI (blue) to visualize actin and nuclei. This representative image corresponds to the 4th to 5th slices of a stack comprising 11 slices (1 being at the bottom). In A, B and D, yellow arrowheads point to TNTs that are visible in the shown pictures, however more TNTs were counted over the whole stack, which were not annotated on the figure. Scale bars correspond to 10 µm. B. Representative images of TNT-connected cells in WT, CD9 KO, CD81 KO and CD9&CD81 KO cells (max projection of 4 to 7 upper slices, z-step 0.4 µm), stained with WGA-488 (green) to label the membrane and DAPI (blue) to label the nuclei. C. Graph of the % of TNT-connected cells in the different cells, from 3 independent experiments. Mean and standard deviation (SD) are: WT = 31.6 ± 1.25; CD9 KO = 25.2 ± 1.24; CD81 KO = 30.4 ± 1.65; CD9 & CD81 KO = 16.3 ± 4.81. p-values are indicated above the brackets in all graphs. D. Representative images (max projection of 5 to 7 upper slices, z-step 0.4 μm) of TNT-connected cells in WT, CD9 OE and CD81 OE cells, stained as in B. E. Graph of the % of TNT-connected cells in the indicated cells from 3 independent experiments. Mean ± SD are: WT = 29.8 ± 1.11; CD9 OE = 45.3 ± 3.17; CD81 OE = 29.5 ± 0.84. F. Vesicle transfer assay from donor cells KO of CD9 and CD81, as schematized on the left. Graphs are % of acceptor cells containing DiD vesicles from 3 independent experiments. Mean ± SD are: WT = 32.7 ± 10.25; CD9 KO = 18.4 ± 8; CD81 KO = 19.1 ± 4.86; CD9 & CD81 KO = 10.5 ± 1.52. G. Vesicle transfer assay from donor cells OE CD9 or CD81, as schematized on the left. Graphs are % of acceptor cells containing DiD vesicles from 3 independent experiments. Mean ± SD are: WT = 29.3 ± 7.45; CD9 OE = 61.3 ± 20.44; CD81 OE = 55.6 ± 10.31.

We then knocked-out (KO) CD9 and/or CD81 by infecting SH-SY5Y cells with lentiviral CRISPR vectors targeting the corresponding genes. Western-blot and Immunofluorescence analysis confirmed the lack of CD9 and/or CD81 expression in these cells (Fig S3A and B). CD9 KO cells, but not CD81 KO cells, showed a significant reduction of the percentage of TNT-connected cells compared to WT cells (Fig 3B&C). The % of TNT-connected cells was even lower in the double KO cells (named CD9&CD81 KO hereafter). The role of CD9 in TNT formation and/or stabilization was confirmed by the finding that CD9 stable overexpression (OE) resulted in a significant increase in the % of TNT-connected cells (Fig 3 D&E). Consistently with KO result, CD81 stable OE did not change the % of TNT-connected cells. Together these results indicated that CD9 plays a role in the formation or maintenance of TNTs and that CD81 might partially complement CD9 absence.

### Positive regulation of TNT function by CD9 and CD81 in donor cells

The next and complementary step was to evaluate the possible influence of CD9 and CD81 on the functionality of TNTs. TNT functionality is understood as the intrinsic capacity to allow the transfer of different types of cellular material through the open channel formed between the cytoplasm of different cells. This was monitored by quantifying by flow cytometry the transfer of labeled vesicles between two different cell populations (“donors” for the cells where vesicles were first labeled and “acceptors” for the cells that received the vesicles [61]). A similar gating strategy was applied to all experiments (Fig S4), and the vesicle transfer through any mechanism other than cell contact-dependent was ruled out in all experiments by analyzing secretion controls, where the two cell populations were cultured separately (see total and secretion transfers in Fig S5). Therefore, the vesicle transfer occurring mainly through cell-contact-dependent mechanisms is an indirect way of monitoring TNT functionality that must be analyzed with regard to TNT apparent number.

First, we co-cultured WT, CD9 KO, CD81 KO or CD9&CD81 KO donor cells versus WT acceptor cells expressing GFP (as schematized in Fig 3F). Consistently with the decrease of the % of cells connected by TNTs, CD9 KO cells showed a significantly decreased percentage of acceptor cells containing donor’s vesicles compared to WT cells (Figs 3F and S4A). On the other hand, despite having no effect on the number of TNTs, CD81 KO resulted in a significant reduction in vesicle transfer to acceptor cells. CD9&CD81 KO cells showed a further decrease of vesicle transfer, consistent with the high decrease in the % of TNT-connected cells. We obtained very similar results when analyzing the experiment using fluorescence microscopy as complementary approach to FACS (Fig S6). Next, we performed a co-culture using tetraspanin OE cells as donor cells (Fig 3G). Consistent with the results of KO cells, CD9 OE significantly increased both the number of TNTs and vesicle transfer compared to WT cells (Figs 3G and S4B) whereas the modest CD81 OE (Fig S3C) stimulated vesicle transfer without significant effect on the number of TNTs (Figs 3E, G and S4B). However, given that the modest OE of CD81 was associated with a decrease in CD9 expression (albeit not statistically significant), an antagonist effect of CD81 on TNTs number would not be detectable using this model.

These data showed that both CD9 and CD81 in donor cells positively regulate the transfer of vesicles through TNTs, which could be in the case of CD9, but not CD81, a direct consequence of an increased number of TNT.

### Pathway of CD9 and CD81 in the regulation of TNTs

To further understand whether CD9 and CD81 have redundant roles or are complementary in the pathway of TNT formation, we knocked-out CD81 in cells over-expressing CD9 (quantification of the total amount of CD9/CD81 in these cells by WB in Fig S3C). This did not prevent the increase of the % of TNT-connected cells observed upon CD9 OE, (Fig 4 A&B), suggesting that CD81 does not play a role in TNT formation when CD9 is present. In contrast to WT cells (Fig 3F), the transfer of DiD-labeled vesicles from CD9 OE cells was no longer sensitive to CD81 KO (Figs 4C and S4C), suggesting that CD9 OE can compensate for the absence of CD81 on vesicle transfer. To determine whether CD81 can compensate for CD9 function in TNT, we knocked-out CD9 in CD81 OE cells (Fig S3C). As shown in Fig 4B, CD9 KO reduced TNT number and vesicle transfer to the same extent in these CD81 OE cells (Figs 4C and S4C) as in WT cells (Fig 3C). Thus, CD81 may not compensate for CD9 for the formation/stabilization of TNTs. This is in accordance with the two tetraspanins having different roles in the process of formation of TNTs.

**Figure 4:**
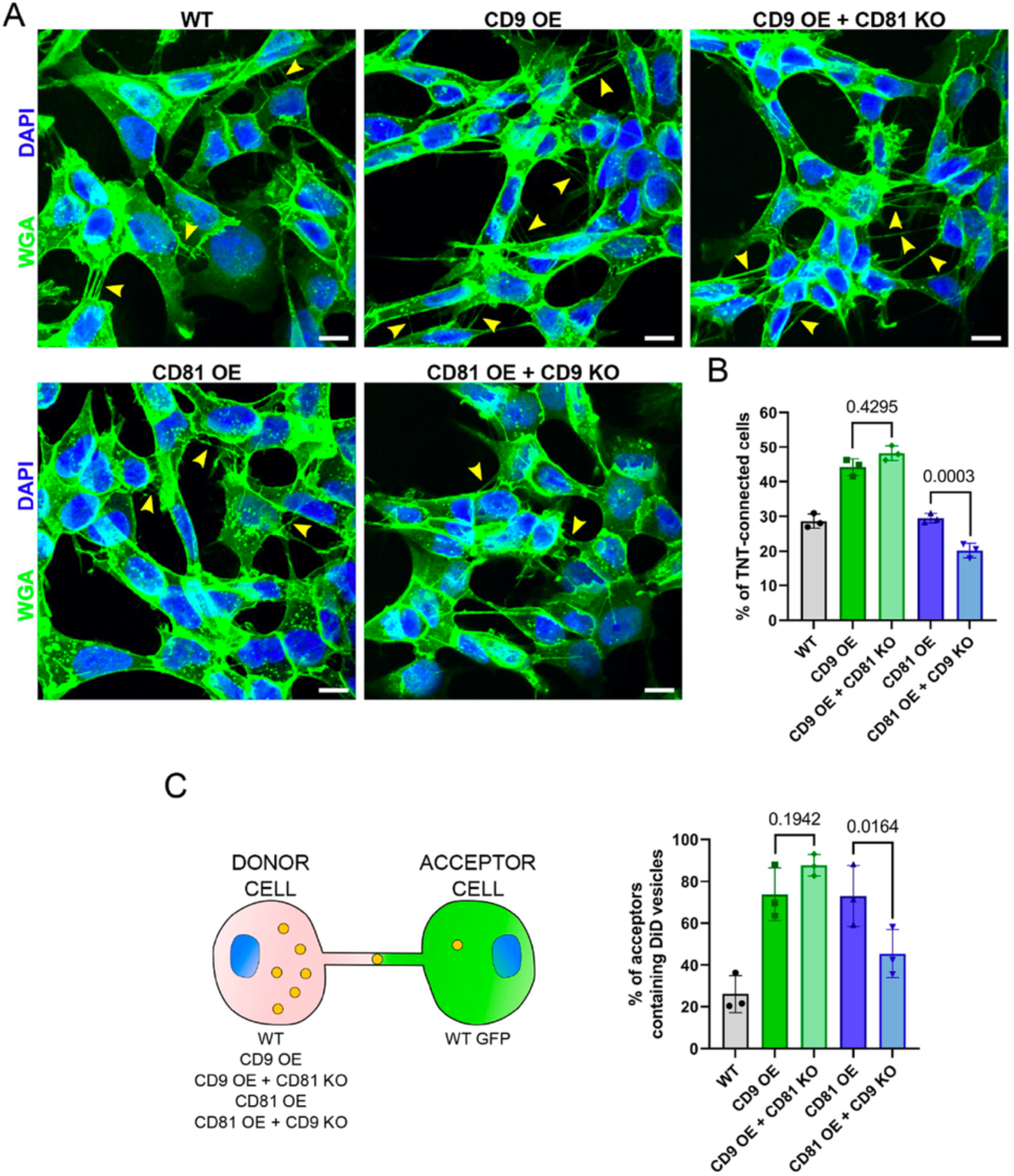
CD9 and CD81 act successively in the formation of TNTs A. Representative images of WT, CD9 OE, CD9 OE + CD81 KO, CD81 OE and CD81 OE + CD9 KO cells, stained with WGA-488 (green) and DAPI (blue). Yellow arrowheads show TNTs. Scale bars correspond to 10 µm. B. Graph of the % of TNT-connected cells in the tetraspanin OE + KO cells. Mean ± SD (N=3) are: WT = 28.6 ± 2.05; CD9 OE = 44.2 ± 2.4; CD9 OE + CD81 KO = 48.2 ± 2.16; CD81 OE = 29.5 ± 1.36; CD81 OE + CD9 KO = 20.2 ± 2.09. p-values are above the brackets. C. Coculture between tetraspanin OE + KO cells used as donors and WT GFP cells used as acceptors, as schematized on the left. The graph shows the % of acceptor cells containing DiD vesicles. Mean ± SD from 3 independent experiments: WT = 26.1 ± 8.94; CD9 OE = 73.8 ± 12.64; CD9 OE + CD81 KO = 87.7 ± 5.11; CD81 OE = 73 ± 14.60; CD81 OE + CD9 KO = 45.4 ± 11.53.

### Stabilization of TNTs by CD9 AB

Knowing that the molecular conformation of CD9 molecules can induce membrane curvature [62,63] and that induced clustering of CD9 with specific antibodies [64,65] could lead to the formation of protrusions such as microvilli [66], we postulated that CD9 could be involved in the initial steps of TNT formation. Consequently, we addressed whether promoting CD9 clustering could affect TNT number and function. Incubation for 2-3 hours of living WT SH-SY5Y cells with an anti-CD9 monoclonal antibody (CD9 AB), but not an antibody targeting another cell surface protein (CD46 AB) or a non-specific control antibody (CTR AB), caused the relocalization of CD9 and CD81 in patches on the plasma membrane, suggesting that these molecules were incorporated in multimolecular complexes that were clustered together by the anti-CD9 antibodies (Figs 5A and S7). Furthermore, CD9 AB treatment led to an increase of more than 30% in the percentage of TNT-connected cells (Figs 5B and S7A). As a control, neither the CD46 AB nor the CTR AB, changed the % of TNT-connected cells (Figs 5A, B and S7A). The result was similar whether the antibodies used on living cells were directly coupled to a fluorophore (Fig 5A, B) or not (Fig S7A), thus eliminating any post-fixation artifact. Addition of CD9 AB after 24-hour coculture of WT donor and acceptor cells for an additional 2 hours resulted in an increase of the vesicle transfer of 30% (Fig 5C). This increase is highly significant considering that it results only from the additional 2-hours treatment over a 24-hours culture compared to control conditions. These data suggested that CD9-enriched sites could serve as initiation platforms for TNTs or participate in the stability of these structures, resulting in increased functionality. The lack of CD81 (using CD81 KO cells) did not impact CD9 clustering (Figs 5D and S7B) or the increase of TNT number induced by the CD9 AB (Figs 5E and S7B). However, it prevented the increase in vesicle transfer (Fig 5F) stimulated by the CD9 AB. This further suggests that CD9 does not require CD81 to form/stabilize TNTs and that CD81 rather regulates TNT functionality.

**Figure 5:**
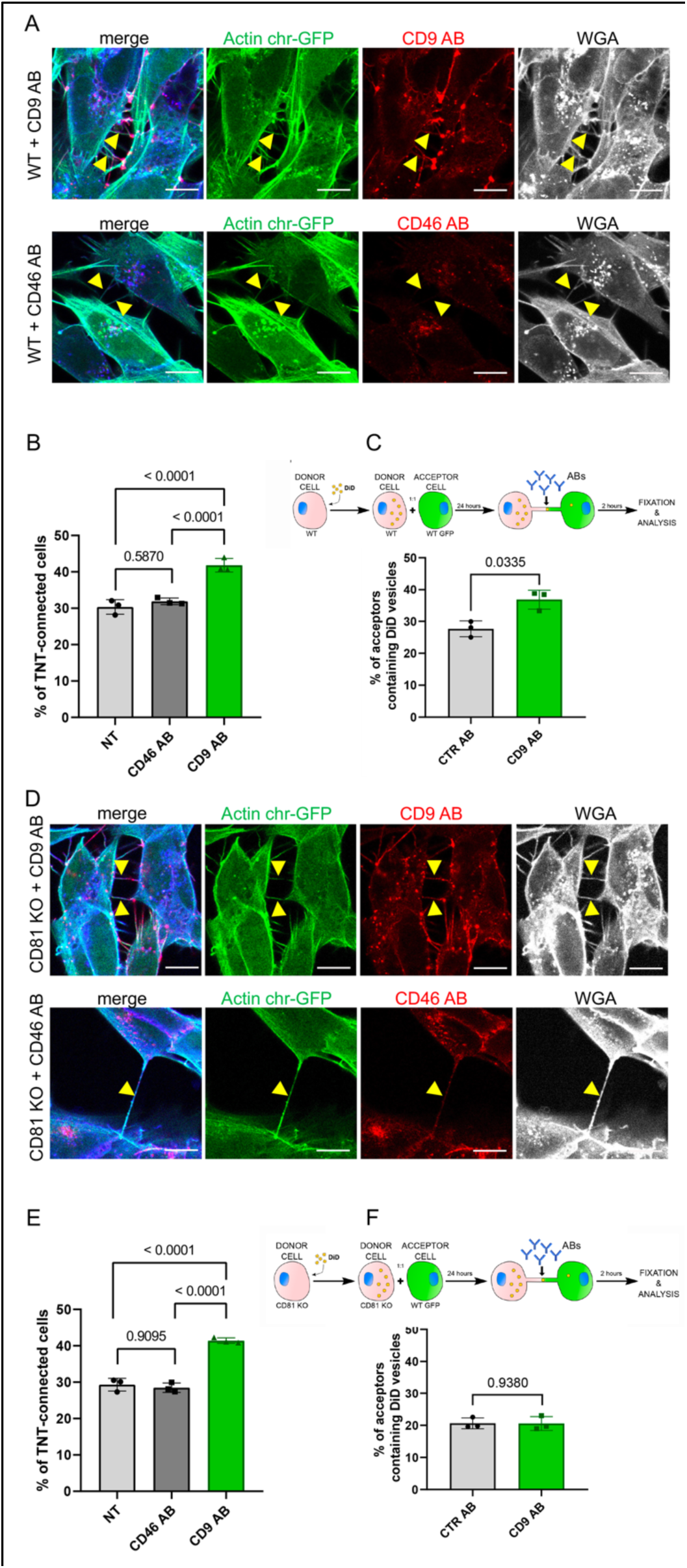
CD9 and CD46 antibody treatment in WT and CD81 KO cells A. Representative confocal images of Actin chromobody-GFP expressing WT cells, treated for 3 hours with either anti-CD9 antibody coupled to Alexa Fluor 568 (upper row), or with anti-CD46 antibody coupled to to Alexa Fluor 594 (second row). Alexa-647 coupled WGA and DAPI were added post-fixation. The images correspond to a maximal projection of 3 or 4 slices of the z-stack encompassing the TNTs indicated with yellow arrowheads. The scale bars correspond to 10 µm. B. Graph of the % of TNT-connected cells in NT, CD46 AB or CD9 AB treatment in WT cells. Mean percentages ± SD (N=3) are 30.4, 31.9 and 41.8, p values resulting from statistical analysis are indicated above the brackets. C. Top, schematic of the antibody treatment experiment after coculture of WT SH-SY5Y donor cells (DiD in yellow circles to stain the vesicles) and WT GFP-labeled acceptor cells, treated with control antibodies (CTR AB) or with antibodies anti-CD9 (CD9 AB) for an additional 2 hours. Graph represents the % of acceptor cells containing DiD vesicles of the cocultures of WT cells with CTR AB or CD9 AB treatment. Mean ± SD (N=3) are: CTR AB = 27.7 ± 2.48; CD9 AB = 36.8 ± 3.03. D. Representative confocal images as in A except that CD81 KO cells were used. E. Graph of the % of TNT-connected cells in NT, CD46 AB or CD9 AB treatment in CD81 KO cells. Mean percentages ± SD (N=3) are 29.3, 28.5 and 41.4, p values resulting from statistical analysis are indicated above the brackets. F. Top, schematic of the antibody treatment experiment after coculture of CD81 KO SH-SY5Y donor cells (DiD in yellow circles to stain the vesicles) and WT GFP-labeled acceptor cells, treated with control antibodies (CTR AB) or with antibodies anti-CD9 (CD9 AB) for an additional 2 hours. Graph represents the % of acceptor cells containing DiD vesicles, mean percentages ± SD (N=3) are: CTR AB = 20.6 ± 1.67; CD9 AB = 20.6 ± 2.16

To determine whether the CD9 AB stabilized TNTs, we monitored TNT duration using time-lapse imaging (as previously set up [8]) of WT or CD81KO SH-SY5Y cells stably expressing actin-chromobody-GFP. As exemplified in movies S1 to S6 and shown in Fig 6A-G, CD9 AB treatment, contrary to CD46 AB or control conditions (without antibody), significantly increased the duration of TNTs from around 11 min to more than 22 min on average, both in WT and CD81KO cells (Fig 6G). This stabilization was accompanied by an enrichment of CD9 at the base of TNTs over time (Fig 6B and E), as observed on fixed cells (Figs 5 and S7), whereas the CD46 AB was internalized and not specifically enriched on TNTs (Fig 6C and F). These results are in accordance with CD9 having a role in stabilizing TNTs, CD81 not being necessary for this step.

**Figure 6:**
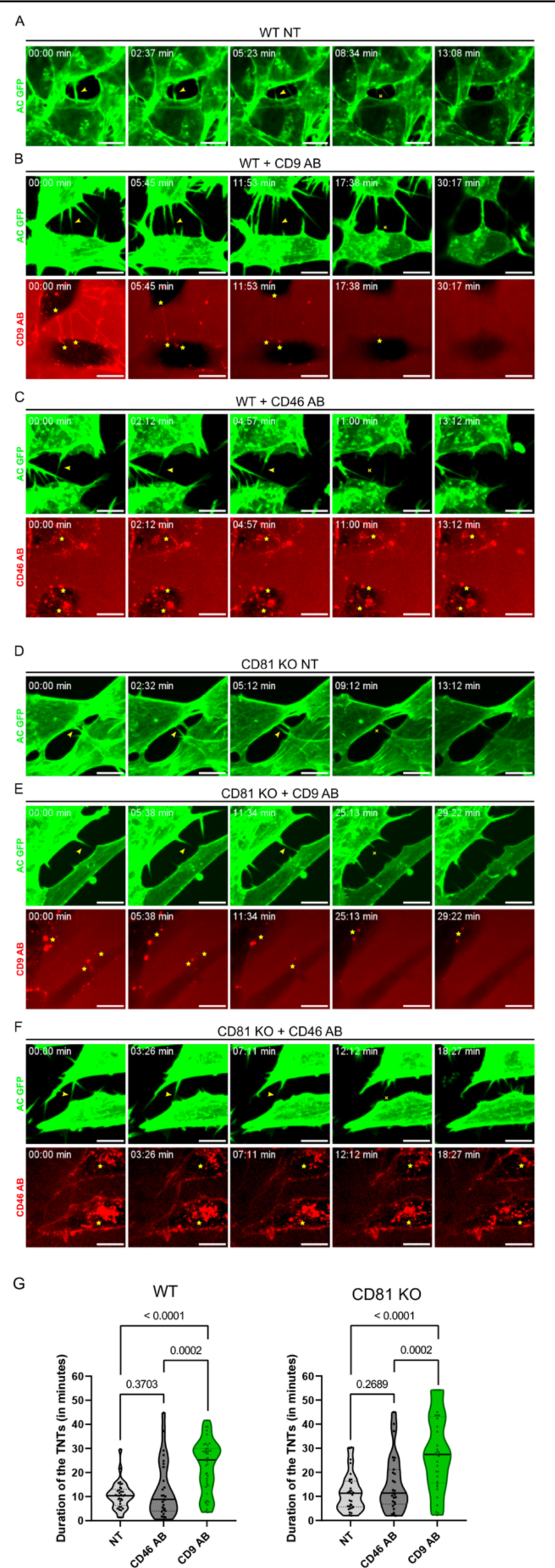
Stabilization of TNTs by CD9 AB. A. Representative snapshots of a TNT over time from Movie S1, corresponding to non-treated WT SH-SY5Y cells (WT NT). In A-F, green panels correspond to the signal of actin chromobody-GFP (AC GFP). Yellow arrowheads in the green panels are pointing to TNTs and the yellow crosses mark the breakage/dissociation of the TNT. Yellow stars in red panels mark the accumulation of CD9 AB (at the base/tip of the TNTs) or CD46 AB (intracellularly). Scale bars correspond to 10 µm. B. Representative snapshots of TNTs over time from Movie S2, corresponding to WT SH-SY5Y cells treated with 10 µg/mL of CD9 specific antibody (WT + CD9 AB). Red panel corresponds to the signal of CD9 antibody coupled to Alexa Fluor 568. C. Representative snapshots of TNTs over time from Movie S3, corresponding to WT SH-SY5Y cells treated with 10 µg/mL of CD46 specific antibody (WT + CD46 AB), coupled to Alexa Fluor 594 (red panel). D. Representative snapshots of a TNT over time from Movie S4, corresponding to non-treated CD81 KO SH-SY5Y cells (CD81 KO NT). E. Representative snapshots of TNTs over time from Movie S5, corresponding to CD81 KO SH-SY5Y cells treated with 10 µg/mL of CD9 specific antibody (CD81 KO + CD9 AB), coupled to Alexa Fluor 568. F. Representative snapshots of TNTs over time from Movie S6, corresponding to CD81 KO SH-SY5Y cells treated with 10 µg/mL of CD46 specific antibody (CD81 KO + CD46 AB) coupled to Alexa Fluor 594. G. Average duration of TNTs in WT (left) or CD81 KO cells (right), measured by live imaging from 3 independent experiments, and represented in Violin plots (with line at median). Left: mean lifetime of 33 TNTs in WT non-treated (NT) cells was 10.5 minutes (± 5.59), 12.8 minutes (± 11.68) in 27 TNTs measured for WT cells treated with CD46 AB, while for WT cells treated with CD9 AB the average duration of the 40 TNTs measured was 22.6 minutes (± 10.87). Right: mean lifetime of TNTs in CD81 KO non-treated (NT) cells, CD81 KO cells treated with CD46 AB, and CD81 KO cells treated with CD9 AB. The average lifetime of 31 TNTs in CD81 KO NT cells was 11.7 minutes (± 7.78), 15.3 minutes (± 11.70) in 27 TNTs in CD81 KO cells treated with CD46 AB, while the average duration of 33 TNTs in CD81 KO cells treated with CD9 AB was also 28.2 minutes (± 15.66). Statistical analysis was performed using one-way Anova with Holm-Sidak’s multicomparison test.

### Regulation of TNT completion by CD81

We next tried to identify the mechanism leading to the reduced vesicle transfer from CD81 KO cells. One possibility being that vesicles cannot enter these connections in the absence of CD81. Timelapse microscopy was used to monitor the impact of CD81 KO on the behavior of DiD-positive vesicles in TNTs. Movies S7 shows an example of vesicle entering TNTs from WT cells and reaching the neighboring cell with constant speed and direction. Movies S8, S9 show examples of vesicles from CD81 KO cells getting stuck inside TNTs or at the basis of TNTs. We quantified the content of TNTs in DiD-positive vesicles by fixing the cells 8 or 24 hours after cell labeling with DiD. Fig 7A shows that the proportion of TNTs containing DiD-labelled vesicles (examples in the left-hand panels) was significantly increased in CD81 KO cells compared to WT cells. Thus, the decrease in transfer observed in the absence of CD81 (Fig. 3F) is not due to a decrease in vesicle entry into TNTs, but rather to the fact that vesicles enter TNTs but are unable to move the full length and, consequently, accumulate to some extent in the TNTs. Two hypotheses could explain this result: i) a decreased fusion of TNT with acceptor cells or ii) an incomplete formation of individual iTNTs in the TNT bundle (iTNTs that would not reach the opposite cell). The latter hypothesis takes into account the fact that the TNTs of SH-SY5Y cells are formed by closely apposed bundles of iTNTs which could originate by opposite cells [2] and are detected as one single TNT by confocal microscopy. Thus, CD81 KO would have an impact on TNT functionality, but not on its detection as a whole structure by confocal microscopy if TNT are formed by iTNTs originating from the two cells. To address this hypothesis, we performed cocultures (Fig 7B) of cells expressing Actin chromobody-GFP (considered as donor cells) with WT cells (acceptor cells). After fixation, actin and cell membranes were stained with phalloidin and WGA respectively. TNTs were identified as phalloidin (which labels all TNTs and iTNTs, no matter the cell of origin) and WGA-positive structures, whereas green fluorescence only labeled actin of iTNTs growing from donor cells, as schematized in Fig 7B. After max projection of the z slices encompassing the TNTs of interest, we classified the TNTs into three categories based on their actin chromobody-GFP content: 1. overlapping with phalloidin throughout the structure (fully green exemplified in the first row of Fig 7B pictures); 2. partially overlapping with phalloidin or 3. not present at all in the TNTs (exemplified in second and third rows respectively). These different possibilities (summarized in the diagram of Fig 7B) could respectively correspond to TNTs whose iTNTs from donor cells reached the opposite cells (1), did not reach acceptor cells or were non-open (2), and to TNTs growing only from acceptor cells (which were WT, (3)). As shown in Fig 7C graph, the percent of fully green TNT dropped from 69% in experiments with WT donor cells to 34% using CD81 KO donor cells, at the profit of partially green TNT, “not green TNT” being minor in both cultures. These results suggested that in CD81KO cells, iTNT growth could be induced, but less iTNTs grow all the way to the acceptor cells, resulting in less transfer to acceptor cells. These results are consistent with CD81 favoring the full achievement of TNT formation, possibly playing a role in iTNT membrane docking to or fusion with opposing membrane whereas CD9 would stabilize TNT (see working model in Fig 7D).

**Figure 7:**
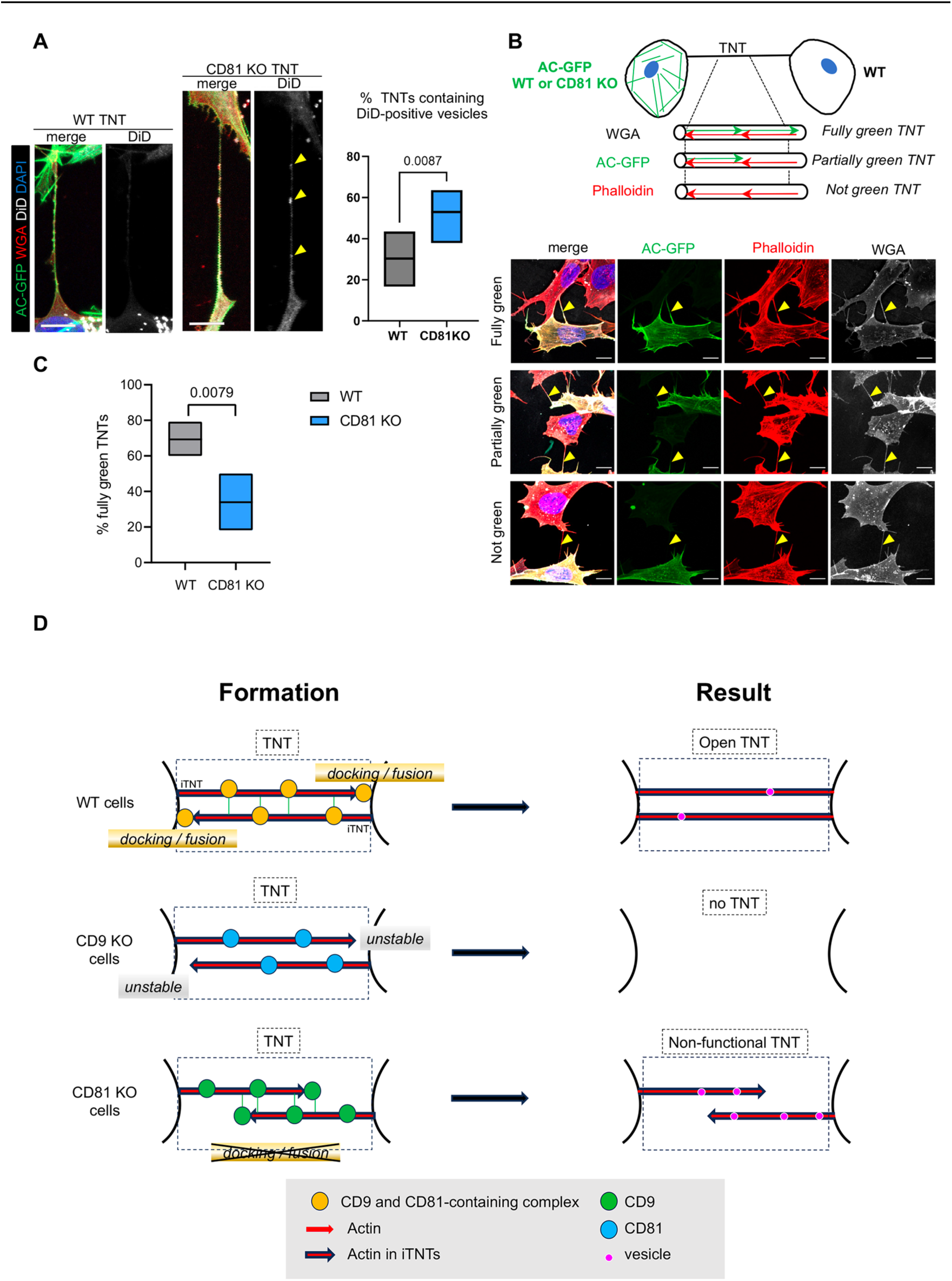
Completion of TNTs by CD81 A. DiD vesicles in TNTs from WT or CD81 KO cells. Actin chromobody-expressing SH-SY5Y cells, either WT or CD81 KO, were challenged with DiD for 30 min and fixed to preserve TNTs after 8 or 24 hours. After additional WGA staining, TNTs (above 10 µm in length) were identified by confocal microscopy using WGA channel, and classified according to their content in DiD vesicles. Left is shown representative examples of long TNTs containing (projection of 12 slices of 0.19 µm) or not (projection of 7 slices) DiD-positive vesicles, from CD81 KO and WT cells respectively. Scale bars are 10 µm, yellow arrowheads point to DiD-positive vesicles in the TNT. The floating bar graph on the right shows the percentage of TNTs that contain DiD-positive vesicles from 6 independent experiments (min to max results, line at means (30.3 and 53.0), medians are 33.4 and 53.7. Statistical analysis is Mann-Whitney test, p-value is above the brackets. TNT number per experiment: 21, 23, 19 (8h of treatment), 6, 30, 41 (24h of treatment) for WT cells; 22, 17, 29 (8h of treatment), 17, 22, 29 (24h of treatment) for CD81KO cells. See also movies S7, S8, S9. B. Coculture of actin chromobody-GFP-expressing cells (WT or CD81 KO) with WT cells. After fixation, cells were additionally labeled with WGA (Alexa 647, in white) and Phalloidin-Rhodamin (red). TNTs were identified by confocal imaging and classified as schematized above the diagrams: fully green when red and green signals overlapped throughout the structure (exemplified in the first row of pictures), partially or not green when green signal was interrupted or not present at all in the TNTs (exemplified in second and third rows respectively). First and third rows are maximal projections of 6 and 7 slices (of 0.19 µm) encompassing the TNT of WT-WT coculture, second row is a maximal projection of 5 slices of CD81 KO-WT coculture. Yellow arrowheads point to TNTs. Scale bars are 10 µm. C. Quantification of the percentage of fully green TNTs in the cocultures of panel B, from 5 independent experiments (approximately 20 TNTs per condition and experiment, in total 91 and 92 analyzed TNTs from WT-WT and CD81 KO-WT cocultures respectively). Floating bars are min to max results, line at means (69.3 and 34%), medians are 66.7 and 27.8 %, statistical analysis was Mann-Whitney test. D. Working model of CD9/CD81 roles on TNT formation. TNTs (black dotted-line frame) are made up of iTNTs, supported by actin (red arrows). In WT cells, CD9 and CD81 are present on growing iTNT (probably together with additional interactors, indicated as yellow circle), and CD9 stabilizes them. At the junction between iTNT and opposing cell membrane, CD81 participates in membrane docking/ fusion, which results in the full completion of an open, functional TNT (right panel). When cells are treated with CD9AB, the active complexes are further enriched/stabilized, and more TNTs are stabilized. In CD9 KO cells, growing iTNTs are not stabilized, resulting in the decrease of TNT number. In CD81 KO cells, CD9-containing complexes (green circles) are still active for stabilization of iTNTs but anchoring of the iTNTs to opposite cell/fusion is no longer induced by CD81, resulting in apparent, but non-functional TNTs, in which vesicles (purple circles) are trapped.

## Discussion

### The composition of TNTs is unique

Thanks to a newly established procedure in U2OS cells, we were able to reproducibly isolate a cellular fraction from cell bodies and from EVPs, and to analyze their content by mass spectrometry. The physical characteristics as well as the proteome analysis of this fraction, which is the TNT-microsomal-type/membrane proteome (called TNTome for simplicity) suggested that it could be enriched in TNTs with minor contaminations with other cell protrusions material. Therefore, its qualitative analysis could give valuable information on core components, traveling material or regulatory factors of TNTs, although we cannot rule out that some are specific to U2OS cells. Our results regarding the unique composition of TNTs were in accordance with those obtained by Gousset et al. [24], who used a laser captured microdissection approach combined with mass spectrometry to reveal the composition of various types of cellular protrusions in mouse CAD cells, including growth cones, filopodia and TNTs. Interestingly, among the 190 proteins identified in [24] in 2 samples from hCAD samples (enriched in TNTs), 101 were also found in TNTome (listed in table S7), mostly corresponding to the most abundant cellular proteins. It is possible that compared to our approach, laser microdissection allowed a limited amount of material to be purified, and therefore few membrane proteins were identified.

TNTome is rich in membrane-associated proteins, as well as in proteins linked to the cytoskeleton, in particular to actin, as it was expected for these structures. TNTome is also abundant in ribonucleoproteins and translation-related factors (220 proteins, 19%). Some of them (as well as the proteins identified as nuclear in the databases) could be contaminants coming from cellular debris, however this fraction should not be more than 10% of the total, based on nanoFCM results. Together with the fact that TNTs transfer mitochondria and lysosomes, it is possible that TNTs are used as a route to transfer RNAs alone or tethered to organelles by Annexins (several members in TNTome, including A11), G3BP1 or CAPRIN1 (both in the first quartile of TNTome [67–69]) or that local translation happens along the TNT to fuel it with required material. Transfer of mRNAs and microRNAs through TNTs has been described [49,50] and could be one of the major functions of this kind of communication.

Regarding membrane, beside tetraspanins, cadherins and integrins, some other plasma membrane TM proteins are of interest. Several amino acids and monocarboxylate transporters (respectively, SLC3A2, SLC1A5 in the first quartile, SLC1A4 and SLC7A1 in the fourth; and SLC16A1 in the second quartile) were found, suggesting together with the presence of mitochondria and of many metabolic enzymes that active metabolism takes place in TNTs. This may be necessary to generate local ATP to power TNT growth as previously suggested [70]. Also, of great interest are SLC1A4 (neutral amino acid transporter) and STX7, which are the only TM proteins uniquely found in TNT and not EVP fraction. Overall, our results suggest that TNTs are different from EVPs, although they share numerous factors. Further studying TNT proteins not present in EVPs will possibly allow to identify TNT specific markers.

To further validate our proteomic approach, we focused our attention on tetraspanins CD9 and CD81. Among the structures described to comprise at least one of these factors and that could potentially be contaminants of our TNT preparations, are midbodies. Midbodies have been shown to contain CD9 and its partners CD9-P1 and EWI-2 [71], and these structures could be copurified with TNTs, although in lower amounts possibly because cells were maintained in serum-free medium for 24 hours before harvesting TNTs. Of note, only 13 TM proteins are common between TNTome and flemingsome (which has 29), including CD9, CD9P1, EWI2, but also CADHF1 and 4, CD44, Integrin α3. Some of these proteins could have a specific role in both TNTs and midbodies, like CD9 and its associated factors. Alternatively, some of these proteins, relatively abundant on plasma membranes, could be randomly present, independently of the specific structure that is analyzed. Importantly, no other specific markers of midbodies were detected in our MS data, including CRIK, CEP55, PLK1, PRC1, MKLP1 and 2, although they are expressed in U2OS cells [56,57] confirming that TNTs and midbodies have different composition.

Among the cellular structures that depend on tetraspanin-enriched microdomains are also the recently described migrasomes, which are substrate-attached membrane elongated organelles formed on the branch points or the tips of retraction fibers of migrating cells which allow the release and transfer of cellular material in other cells [72]. Although migrasomes have been described to be enriched in, and dependent on tetraspanins-enriched microdomains [73,74], they are fundamentally different from TNTs since they are attached to the substratum (and probably not collected during the procedure of TNT purification), and exhibit specific markers, identified by MS analysis, which are absent from TNTome (TSPAN 4 and 9, NDST1, PIGK, CPQ, EOGT for example, see [75]). Altogether, our TNT purification protocol and proteomic analysis have revealed that TNTs have specific composition opening new avenues to understand how TNTs are formed and regulated.

### CD9 and CD81 regulate formation of TNTs

TNTome in U2OS revealed that CD9 and CD81 are among the most abundant integral membrane proteins in TNTs (see table S3, tab2). Consistent with previous results of overexpression [26], our data showed that they are also present in TNTs in other cell lines, especially SH-SY5Y cells, making them probable compulsory proteins of TNTs. Based on previously reported roles for these proteins, CD9 and CD81 were interesting candidates to play a role in TNT formation and/or function. Indeed, CD9 senses membrane curvature [76], and several studies have reported a strong impact of these tetraspanins on membrane extensions [62], the most striking example being a profound modification of microvilli shape and distribution at the surface of CD9 KO oocytes [77]. This may be due to their inverted cone-like structure which can induce membrane deformation in vitro [63]. In addition, both CD9 and CD81 have been shown to be involved in fusion, an important step in TNT formation following membrane extension; as positive regulators of sperm-egg fusion, but negative regulators of macrophage and muscle cell fusion [36,78–81]. We show here that CD9 and CD81 are positive regulators of TNT function, since cell contact-dependent vesicle transfer in coculture experiments is decreased when CD9 or CD81 is lacking. Interestingly and despite their abundancy in EVPs, CD9 and/or CD81 absence did not affect transfer through secretion. These results confirmed recent data from the literature showing that CD9 and CD81 have minimal impact on the production or composition of EVs in MCF7 cells [82] and that the lack of CD9 does not impair the delivery of the content of EVs into recipient cells [83]. Interestingly, EVs have been shown to regulate TNT formation in mesothelioma, breast cancer or brain endothelial cell models [84–86]. Although we cannot completely rule out a crosstalk between EVP and TNT formation or stability, our results together with the current data in literature indicate that CD9 and CD81 deficiencies have a stronger impact on TNT function than on EVP production or function, suggesting that EVPs and TNTs are regulated independently of each other, as far as tetraspanins are concerned.

### Although CD9 and CD81 can partially compensate for each other, they act at different steps of TNT formation

Overall, our results indicate that CD9 and CD81 regulate cell-contact-mediated transfer, through different mechanisms. CD9 positively regulates the % of cells connected by TNTs, likely by stabilizing the TNTs. Indeed, short-term CD9 antibody treatment significantly increased TNT-connected cells and vesicle transfer, and this was associated with an increased TNT duration. CD9 antibody treatment also relocated CD9 and CD81 to TNT extremities, supporting a role of CD9-containing complexes in stabilizing TNTs maybe by favoring cis and/or trans interactions.

In contrast to CD9, CD81 regulates vesicle transfer without any effect on the number of TNT-connected cells, suggesting that these TNTs are less functional. Compared to WT cells, live cell imaging of CD81 KO cells showed that vesicles can still enter TNTs but do not efficiently transfer to the opposite cells, therefore accumulating to some extent inside TNTs, as confirmed by the increased number of TNTs that contain vesicles after fixation. One possible explanation could be a reduced ability of CD81 KO TNTs to fuse with the opposing cell. However, we have shown that a GFP-labeled actin chromobody reached the opposing cell twice less frequently when expressed in CD81 KO cells than when expressed in WT cells, pointing again to an abnormal structure of these TNTs in the absence of CD81. The TNTs of SH-SY5Y cells are formed by a bundle of closely apposed individual tubes, iTNTs [2], originating from the two connected cells, which by confocal microscopy are detected as one single TNT, due to low resolution. The diminished percentage of TNTs fully labeled by the actin chromobody could therefore be explained by a decrease of the fraction of iTNTs originating from CD81 KO cells that reaches or is stabilized at the opposing cell. Therefore, CD81 could act to facilitate anchoring of the growing iTNT membrane to the opposite membrane, ultimately enabling fusion and completion of fully open/functional TNTs.

Although CD81 does not seem to regulate the number of TNTs, several lines of evidence indicate that these two tetraspanins can compensate for one another in the formation/maintenance of TNTs and their function. First the double KO of CD9 and CD81 reduced TNT-connected cells to levels lower than the CD9 single KO, indicating partially redundant roles for CD9 and CD81 in TNT biogenesis or stability. CD9 can also compensate for the lack of CD81 as its OE bypasses the requirement for CD81 for full vesicle transfer. This is reminiscent of the partial compensation by CD81 of the absence of CD9 on eggs during sperm-egg fusion [78–80]. In contrast, KO of CD9 on CD81 OE cells resulted in a significant decrease in the % of TNT-connected cells and vesicle transfer compared to CD81 OE alone. Furthermore, anti-CD9 antibody treatment of CD81 KO cells increased TNT number and duration to levels seen in WT cells, indicating that CD9 TNT-formation/stabilization capacity was independent from CD81. However, the same antibody incubation stimulate vesicle transfer in WT but not CD81 KO cells, suggesting that CD81 controls a different step. Based on these results, we propose the working model shown in Figure 7D, in which CD9 would act by stabilizing the interactions between iTNTs and bringing CD81 to the docking site between iTNT tip and opposite cell, allowing CD81 to participate in the anchoring of opposite membranes and finally in the opening of the channel. Recent work [74] proposed a role for TEM in forming a rigid ring impairing membrane damage to spread. Similarly, the specific CD9/CD81 TEM could somehow protect the membrane around the TNT site where fusion with the opposite cell occurs. More work in identifying specific TEM composition in TNTs would be needed to address this hypothesis.

In summary, in this study we have shown that TNTs and EVPs are two cellular structures with partially overlapping composition, and that despite being part of both TNTs and EVPs, the tetraspanins CD9 and CD81 are fundamental regulators of TNT formation with complementary roles in the whole process of biogenesis of these structures. TNTome further analysis will be of great help to identify additional proteins, as part of the TEM or not, that could participate in TNT formation and regulation.

## Materials and Methods

### Cell lines, lentiviral preparations, plasmids and transfection procedures

U2OS cells were cultured at 37 °C in 5% CO2 in Dulbecco’s Modified Eagle’s Medium (DMEM + Glutamax, + 4.5 g/l Glucose, + Pyruvate, Gibco), plus 10% fetal calf serum (FCS) and 1% penicillin/streptomycin (P/S). U2OS stably expressing H2B-GFP (Addgene 11680) and actin chromobody GFP (pAC-TagGFP from Chromotek) were obtained by transfection with Fugene HD according to manufacturer’s instructions, followed by sorting of GFP-positive cells. GFP-CD9 expressing U2OS cells were obtained by lentiviral transduction as below, followed by limiting dilution to obtain a clone. SH-SY5Y human neuroblastoma cells were cultured at 37 °C in 5% CO2 in RPMI-1640 (Euroclone), plus 10% FCS and 1% P/S.

For the lentiviral preparations, human HEK 293T cells were cultured at 37 °C in 5% CO2 in DMEM (ThermoFisher), with 10% FCS and 1% P/S. Cells were plated one day before transfection at a confluency around 70%. Transfection of the different plasmids was made in a ratio 4:1:4 using lentiviral components pCMVR8,74 (Gag-Pol-Hiv1) and pMDG2 (VSV-G) vectors and the plasmid of interest respectively using FuGENE HD (Promega) according to manufacturer’s protocol. CRISPR lentiviral plasmids containing the sequences for CD9: GAATCGGAGCCATAGTCCAA and CD81: AGGAATCCCAGTGCCTGCTG were selected using the CRISPR design tool available at the Broad Institute (https://portals.broadinstitute.org/gpp/public/analysis-tools/sgrna-design). The corresponding guide DNA sequences were cloned into the lentiCRISPRv2 plasmid (#52961; Addgene) according to the instructions of the Zhang laboratory (https://www.addgene.org/52961/). The viral particules were collected and concentrated using LentiX-Concentrator (TakaraBio) after 48 h. To KO CD9 and/or CD81, SH-SY5Y were plated the day before the infection at a confluency of around 70% and next day the lentivirus was added to the cells. 24 hours later the medium with the lentiviruses was removed and cells were selected with RPMI + 1 μg/mL of puromycin for 2 days. After this, cells were splitted 1:5 and reinfected for 24 hours. After these 24 hours, medium was replaced with RPMI + 1 μg/mL of puromycin for 10 days changing the medium every 2-3 days. Finally, cells were tested for the absence of expression of CD9 and/or CD81 by western blot (WB). These cells were used to generate Actin chromobody-GFP expressing cells, by transfection of the plasmid (pAC-TagGFP from Chromotek) using Lipofectamine 2000 according to manufacturer’s instructions, next sorted to enrich in GFP-positive cells, which were finally cloned by limiting dilution.

To obtain clones that overexpress CD9 or CD81, SH-SY5Y were plated the day before at a confluency of around 70% and next day cells were transfected with the corresponding plasmid using Lipofectamine 2000 (Invitrogen) following the manufacture recommendations and 72 hours post-transfection cells selected with 350 μg/mL of Hygromicin B (Gibco) for 5 days, changing the medium every 2 days and then with 50 μg/mL of Hygromicin B for 10 days more. The CD9 or CD81 OE clones were obtained by limiting dilution and tested for expression of CD9 or CD81 by immunofluorescence and WB.

### TNT and EVP preparation

Two million U2OS cells were plated in 75 cm^2^ flasks for 24 hours (8 flasks per point), next complete medium was replaced by medium without FCS for an additional 24 hours. When necessary, the medium was supplemented with 100 nM Latrunculin A (Millipore). For EVP preparation, conditioned medium was collected, centrifuged twice at 2000g to remove cells, concentrated 10-fold on Vivaspin 20 (MWCO 10kD, Cytiva), and next submitted to ultracentrifugation in a Beckman MLS50 rotor at 10,000g for 30 minutes at 4°C. Supernatant was collected and centrifuged at 100,000g for 70 minutes at 4°C, resulting pellet was resuspended in PBS and centrifuged again at 100,000g for 70 minutes at 4°C. Pellets were used for electronic microscopy, Mass Spectrometry, or solubilized in 2x Laemmli for WB analysis.

For TNT preparation, cell cultures after removing conditioned medium were washed carefully with PBS, next 2 ml of PBS was added in each flask, which was left on an oscillating shaker for 5 minutes before being shaked (30 sec horizontally, and 4 times 30 sec by banging them vigorously). PBS from all flasks was drained, collected and centrifuged twice at 2,000g, and filtered on 0.45 μM syringe filter (Corning) to remove detached cells, next submitted to ultracentrifugation at 100,000g for 70 minutes at 4°C. Pellets were used for electronic microscopy, Mass Spectrometry or WB. After collecting EVPs and TNTs from cell cultures, cells were harvested in PBS, and cell extracts were prepared in Tris 50 mM pH 7.4, NaCl 300 mM, MgCl2 5 mM, Triton 1% with protease inhibitors (complete mini, Roche).

### Negative staining and Transmission Electron Microscopy

EVP or TNT pellets were resuspended in 50 ml of PBS. Four microliters of each sample were spotted on a carbon-coated grid primarily glow discharged and incubated at room temperature for 1 min. Uranyl acetate (2%) in water was used to contrast the grids and incubated for 1 min. The grid was then dried and observed under 120 kV using a Tecnai microscope (Thermo Fisher Scientific) and imaged using a 4000 by 4000 Eagle camera (Thermo Fisher Scientific). To quantify EVP diameter, vesicles were segmented manually using the Quick selection tool of PHOTOSHOP v23.5.5 (Adobe Systems, San Jose, CA). Spot detector under ICY software (https://icy.bioimageanalysis.org/) saved the surface area of the segmented vesicles to a .excel file. The vesicle size d was calculated from the surface area A using d = √4A/π, thereby assuming that vesicles are spherical. For TNTs, ROIs were defined under ICY, and first and second diameters were used for diameter and length respectively.

### Nano-Flow Cytometry

The size and number of exosomes and TNT particles were identified by Nano-Flow Cytometry (NanoFCM). NanoFCM is applicable when the refractive index of input samples is the same or similar to that of silica particles. The standard working curve of scattering light intensity was established using silica standard sphere. EVPs and TNT particles were isolated from 1 flask of culture following the protocol above except the ultracentrifugation steps. The particle size distribution of samples was measured based on the scattering intensity.

### Mass spectrometry

#### Digestion of TNT and EVPs samples

Protein pellets were dissolved in urea 8M, Tris 50mM pH 8.0, TCEP (tris(2-carboxyethyl)phosphine) 5mM and SDS (Sodium Dodecyl Sulfate) 2%. SDS was removed using a methanol/ chloroform/ water extraction. Briefly, 3V of ice-cold Methanol was added to the sample then mixed. 2 V of ice-cold chloroform was added and mixed. Then 3Vof ice cold water was added and mix. Sample were spinned for 3 min at 5000g at 4°C. Proteins at the organic/inorganic interface were kept and washed 3 times in ice cold methanol. Protein pellet were dissolved in Guanidine 1M, TCEP 5mM, Chloroacetamide 20mM, Tris 50mM pH8.0 and samples were heated 5 min at 90°C before digestion in a mix of 500 ng of LysC (Promega) at 37°C for 2h. Dilution with Tris 50mM was done before the addition of trypsin 500 ng (Promega) and digestion at 37°C for 8h. Digestion was stopped by adding 1% final of formic acid. Peptides were purified using a C18 based clean up standard protocol done using Bravo AssayMap device.

#### LC-MS/MS analysis of TNT and EVPs

LC-MS/SM analysis of digested peptides was performed on an Orbitrap Q Exactive Plus mass spectrometer (Thermo Fisher Scientific, Bremen) coupled to an EASY-nLC 1200 (Thermo Fisher Scientific). A home-made column was used for peptide separation (C_18_ 50 cm capillary column picotip silica emitter tip (75 μm diameter filled with 1.9 μm Reprosil-Pur Basic C_18_-HD resin, (Dr. Maisch GmbH, Ammerbuch-Entringen, Germany)). It was equilibrated and peptide were loaded in solvent A (0.1 % FA) at 900 bars. Peptides were separated at 250 nl.min^-1^. Peptides were eluted using a gradient of solvent B (ACN, 0.1 % FA) from 3% to 22% in 140 min, 22% to 42% in 61 min, 42% to 60% in 15 min (total length of the chromatographic run was 240 min including high ACN level step and column regeneration). Mass spectra were acquired in data-dependent acquisition mode with the XCalibur 2.2 software (Thermo Fisher Scientific, Bremen) with automatic switching between MS and MS/MS scans using a top 10 method. MS spectra were acquired at a resolution of 70000 (at *m/z* 400) with a target value of 3 × 10^6^ ions. The scan range was limited from 400 to 1700 *m/z*. Peptide fragmentation was performed using higher-energy collision dissociation (HCD) with the energy set at 26 NCE. Intensity threshold for ions selection was set at 1 × 10^6^ ions with charge exclusion of z = 1 and z > 7. The MS/MS spectra were acquired at a resolution of 17500 (at *m/z* 400). Isolation window was set at 2.0 Th. Dynamic exclusion was employed within 35 s.

All Data were searched using MaxQuant (version 1.6.6.0) using the Andromeda search engine [87] against a human reference proteome (75088 entries, downloaded from Uniprot the 29^th^ of October 2020).

The following search parameters were applied: carbamidomethylation of cysteines was set as a fixed modification, oxidation of methionine and protein N-terminal acetylation were set as variable modifications. The mass tolerances in MS and MS/MS were set to 5 ppm and 20 ppm respectively. Maximum peptide charge was set to 7 and 5 amino acids were required as minimum peptide length. At least 2 peptides (including 1 unique peptides) were asked to report a protein identification. A false discovery rate of 1% was set up for both protein and peptide levels. iBAQ value was calculated. The match between runs features was allowed for biological replicate only.

#### Data analysis

Quantitative analysis was based on pairwise comparison of protein intensities. Values were log-transformed (log2). Reverse hits and potential contaminant were removed from the analysis. Proteins with at least 2 peptides were kept for further statistics. Intensity values were normalized by median centering within conditions (normalizeD function of the R package DAPAR [88]). Remaining proteins without any iBAQ value in one of both conditions have been considered as proteins quantitatively present in a condition and absent in the other. They have therefore been set aside and considered as differentially abundant proteins. Next, missing values were imputed using the impute.MLE function of the R package imp4p (https://rdrr.io/cran/imp4p/man/imp4p-package.html). Statistical testing was conducted using a limma t-test thanks to the R package limma [89]. An adaptive Benjamini-Hochberg procedure was applied on the resulting p-values thanks to the function adjust.p of R package cp4p [90] using the robust method described in [91] to estimate the proportion of true null hypotheses among the set of statistical tests. The proteins associated to an adjusted p-value inferior to a FDR level of 1% have been considered as significantly differentially abundant proteins.

#### Bioinformatic analysis and data mining

Twelve replicates were done for the discovery of TNT proteins. Nine over 12 were kept as being part of the TNT. The 1177 resulting proteins were sorting by quartiles according to their iBAQ value. For each of the 4 lists, proteins composition was depicted using ProteoMap tool [92] to visualize their weighted GO organization and protein contribution for each GO term. Also, a DAVID analysis [93] was done for each quartile of proteins. Functional charts and cluster were used to describe the dataset. Protein networks were visualized using STRING [94] and Cytoscape (cytoscape.org).

Resource availability: The mass spectrometry proteomics data have been deposited to the ProteomeXchange Consortium via the PRIDE[1] partner repository with the dataset identifier PXD033089.

### Sample preparation for TNT imaging

SH-SY5Y or U2OS-derived cells were trypsinized and counted, and 100,000 or 30,000 cells respectively were plated on coverslips. After O/N culture, cells were fixed with specific fixatives to preserve TNT [61], first with Fixative 1 (2% PFA, 0.05% glutaraldehyde and 0.2 M HEPES in PBS) for 15 min at 37 °C, then with fixative 2 (4% PFA and 0.2 M HEPES in PBS) for 15 min at 37 °C. After fixation, cells were washed with PBS and membranes were stained with conjugated wheat germ agglutinin (WGA)-Alexa Fluor (1:300 in PBS, Invitrogen) or Phalloidin-Texas-red (1:300 in PBS, Invitrogen) and DAPI (1:1000) (Invitrogen) at room temperature 15 minutes. After gently washing 3 times with PBS, samples were mounted on glass slides with Aqua PolyMount (Polysciences, Inc.). Every different SH-SY5Y cell type (WT, tetraspanin KO or tetraspanin OE) was prepared in the exact same conditions.

### Quantification of the percentage of TNT-connected cells (also referred to as TNT counting or TNT number)

For U2OS-derived cells, a mild trypsinization was used to be able to visualize connections (including TNTs) between cells in the same flasks used for TNT preparation. Cells were incubated for 4 minutes at 37°C with a 100-fold dilution of 0.05%Trypsin-EDTA solution (Gibco), next cells were fixed as above, and phase images were acquired using Incucyte system (Essen Bioscience, 20x magnification). For SH-SY5Y cells grown on coverslips, multiple random Z-stack images of different points on the samples were acquired using an inverted laser scanning confocal microscope LSM 700 or 900 (Zeiss) controlled by the Zen software (Zeiss). The images were analyzed according to the morphological criteria of TNTs: structures that connect two distant cells and that are not attached to the substratum. First slices were excluded from the analysis, and only connections in the middle and upper stacks were considered. Cells that have TNTs between them were marked as cells connected by TNTs, and the number of these cells was compared to the total number of cells in the sample, giving the percentage of cells connected by TNTs. For both cell types, the analysis was performed using ICY software (https://icy.bioimageanalysis.org/), by using the “Manual TNT annotation” plugin. In each experiment, at least 200 or more cells were analyzed in every condition. The images were adjusted and processed with the ImageJ (https://imagej.nih.gov/ij/) or Icy softwares.

### Co-culture assay (DiD transfer assay) and flow cytometry analysis

DiD transfer assays have been described elsewhere [55]. Briefly, the co-culture consisted in two distinctly labeled cell populations: a first population of cells (donors) was treated with Vybrant DiD (dialkylcarbocyanine), a lipophilic dye that stains vesicles, at 1:1000 (Thermo Fisher Scientific) in complete medium for 30 min at 37 °C (Life Technologies), the cells were then trypsinized and mixed in a 1:1 ratio with a different cell population (acceptors) of different color to distinguish them from donors (usually expressing GFP) and the co-culture was incubated O/N.

In the case of the co-cultures with KO, OE or OE + KO of CD9 and CD81, 400,000 donor cells were mixed with 400,000 acceptor cells on 6-well plates for analysis by flow cytometry. After O/N culture, cells were trypsinized, passed through a cell strainer to dissociate cellular aggregates and fixed with 2% PFA in PBS. Finally, these cells were passed through the CytoFLEX S Flow Cytometer (Beckman Coulter) under the control of the CytoExpert Acquisition software. The data were analyzed with FlowJo software following a similar strategy for all experiments: first the samples were gated to exclude cellular debris by plotting the area obtained with the side scatter (SSC-A) and the area obtained with the forward scatter (FSC-A) obtaining all the cells in the sample. Second, within this previous gate, the sample was gated to exclude cell doublets, plotting the width obtained with side scatter (SSC-W) and the area obtained with forward scatter (FSC-A) thus obtaining the singlets. Finally, within the singlet gate, the co-culture was gated using GFP and DiD expression, resulting in four quadrants delimiting double-negative, GFP-positive, DiD-positive and double-positive populations. The % of acceptor cells receiving DiD-vesicles was obtained by calculating the percentage of acceptor cells with labeled vesicles out of the total number of acceptor cells.

In the case of the co-culture of the CD9 AB treatment, 50.000 donor cells were co-cultured with 50.000 acceptor cells on coverslips. Results were analyzed by microscopy as described above, and results were obtained by semi-quantitative analysis using ICY software (http://icy.bioimageanalysis.org/) by calculating the percentage of acceptor cells with labeled vesicles out of the total number of acceptor cells. In each experiment, at least 100 recipient cells per condition were counted. Image montages were built afterward in ImageJ software.

In all co-cultures a control of the transfer by secretion was performed. DiD-loaded donor cells were seeded alone (800,000 cells for flow cytometry and 100,000 cells for microscopy) and cultured for 24 hours. Next, the supernatant from these cells was centrifuged and added to the acceptor cells that had been seeded on the previous day under the same conditions as the donors, these acceptor cells were cultured for an additional 24 hours and next fixed and analyzed in the same way as above mentioned.

### CD9 and CD46 antibody (AB) treatments

Both TNT counting and vesicle transfer in the CD9 or CD46 AB treatment were done in the same way as described, with an additional step after 24 hours of coculture consisting in the incubation of the cells with either Goat anti-rabbit Alexa Fluor 305 (Thermofisher ref: A21068-) control antibodies (CTR AB) or anti-CD9 (TS9, coupled to Alexa 568) or anti-CD46 (coupled to Alexa 594) antibodies at a concentration of 10 μg/mL in RPMI medium for 2 or 3 hours. Subsequently, the cells were fixed and submitted to immunofluorescence.

### Live imaging microscopy

Time-lapse microscopy imaging was performed on an inverted spinning disk microscope (Eclipse Ti2 microscope system, Nikon Instruments, Melville, New York, USA) using 60X 1.4 NA CSU oil immersion objective lens and laser illumination 488 and 561, with optical sections of 0.4 μm. During image acquisition, 37°C temperature, 5% CO_2_, O_2_ and humidity were controlled with a small environmental chamber (Okolab). Cells were plated in Ibidi μ-slide with glass bottom. Image processing and movies were realized using ImageJ/Fiji software.

### Immunofluorescence

For immunofluorescence, 100.000 SH-SY5Y cells or 30,000 U2OS cells were seeded on glass coverslips and after O/N culture they were fixed with 4% paraformaldehyde (PFA) for 15 minutes, quenched with 50 mM NH_4_Cl for 10 minutes and blocked in 2% BSA in PBS for 20 minutes. Primary antibodies mouse anti-CD9 IgG1 (TS9), anti-CD81 IgG2a (TS81), anti-ITGB1 (β1-vjf, IgG1) or anti-CD151 (TS151, IgG1) were previously described [95] and were used in 2% BSA in PBS during 1 hour. Other primary antibodies used after permeabilization were in 0.05% saponin, 0.2%-containing PBS: mouse anti-Vinculin (Sigma V9264, 1:500), rabbit anti-paxillin (Santa-Cruz, sc-5574, 1:1000), mouse anti GM130 (BD 610823, 1:1000). After 3 washes of 10 minutes each with PBS, cells were incubated with each corresponding Alexa Fluor-conjugated secondary antibody (Invitrogen) at 1:1000 in 2% BSA in PBS for 1 hour. Specifically, the secondary antibodies used were goat anti-mouse with epitope IgG1 Alexa Fluor 488 for CD9 (Invitrogen ref: A21121) and goat anti-mouse with epitope IgG2a Alexa Fluor 633 for CD81 (Invitrogen ref: A21136). For the experiments showing the actin cytoskeleton, cells were labeled with Phalloidin-Rhodamine (Invitrogen) in the same mix and conditions as the secondary antibodies. Then, cells were washed 3 times of 10 minutes each with PBS, stained with DAPI and mounted on glass slides with Aqua PolyMount (Polysciences, Inc.). Images were acquired with a confocal microscope LSM700 or 900 (Zeiss) and processed with the ImageJ or Icy softwares.

### Western blot

For Western blot SH-SY5Y cells were lysed with lysis buffer composed of 150 mM NaCl, 5 mM EDTA, 20 mM Tris, pH 8.0, 1% Triton X100 with Roche *cOmplete™* Protease Inhibitor Cocktail. Protein concentration was measured by a Bradford protein assay (Bio-Rad). 20 μg of protein were submitted to PAGE (in non-reducing conditions for tetraspanins and ADAM10) followed by western blot analysis. Primary antibodies used for Western blot were: mouse anti-CD9 IgG1 (TS9, 1:1000), mouse anti-CD81 IgG2a (TS81, 1:1000), mouse anti CD63 (1:1000), mouse anti ADAM10 (11G2, 1:1000) [34,95], rabbit anti-α-GAPDH (Sigma ref: G9545, 1:1000), mouse anti-actin (MP Biomedicals, clone C4, 1:1000), mouse anti-α-tubulin (Sigma ref: T9026, 1:2000), mouse anti-GM130 (BD transduction Laboratories, ref:610822, 1:1000, rabbit ITGB1, ITGB4, ITGA4 (from integrin Ab Sampler kit, Cell signaling ref: 4749, 1:1000), rabbit anti-EGFR (Cell Signaling ref: 4267, 1:1000), rabbit anti-CX43 (Sigma ref: C6219, 1:3000), mouse anti-ANXA2 (Proteintech ref: 66035,1:2000), mouse anti-Alix (Biorad MCA2493, 1:1000), rabbit anti-Calnexin (Enzo SPA-860, 1:1000).

### Statistical analysis

Statistical analysis of experiments concerning the TNT counting and the DiD transfer assay are described elsewhere [4]. Briefly, the statistical tests were applied using either a logistic regression model computed using the ‘glm’ function of R software (https://www.R-project.org/) or a mixed effect logistic regression model using the lmer and lmerTest R packages, applying a pairwise comparison test between groups.

All graphs shown in this study have been made with GraphPad Prism version 9. The numerical data used in all figures are included in <u>Data S1</u>.

## Acknowledgements

We acknowledge the Center for Translational Science (CRT)-Cytometry and Biomarkers Unit of Technology and Service (CB UTechS), in particular P.H. Commère, as well as G. Pehau-Arnaudet and A. Mallet from the Ultrastructural Bioimaging (UBI) facility. Thanks to Dr. S. Charrin for helpful discussion, Dr. S. Lebreton, R. Chakraborty for critical reading of the manuscript, and all the members of the UTRAF unit for their support. We are endebted to late Ms Marguerite MICHEL whose bequest to Institut Pasteur has made this project possible. We are grateful for financial support to C.Z. from Institut National du Cancer (PLBIO18-103), Association France Alzheimer (AAP PFA 2021 #6156), Equipe Fondation Recherche Médicale (FRM EQU202103012692), and Agence Nationale de la Recherche (ANR-20-CE13-0032 to C.Z. and ANR-21-CE35-0007 to M.M.). We are grateful for support for equipment from the French Government Programme Investissements d’Avenir France BioImaging (FBI, N° ANR-10-INSB-04-01) and the French gouvernement (Agence Nationale de la Recherche) Investissement d’Avenir programme, Laboratoire d’Excellence “Integrative Biology of Emerging Infectious Diseases” (ANR-10-LABX-62-IBEID).

## Author contributions

Conceptualization: RNM, CB, CZ; methodology: RNM, TC, ER, EP, MM, CB; formal analysis: RNM, TC, EP, CB; writing—original draft: RNM, CB; writing— review and editing: CZ, RNM, CB, ER, TC, MM; funding acquisition: CZ, MM; supervision: CB, CZ.

