## Supplementary material for "Proteomic landscape of tunneling nanotubes reveals CD9 and CD81 tetraspanins as key regulators": Suppl fig and legends.

### Figure S1

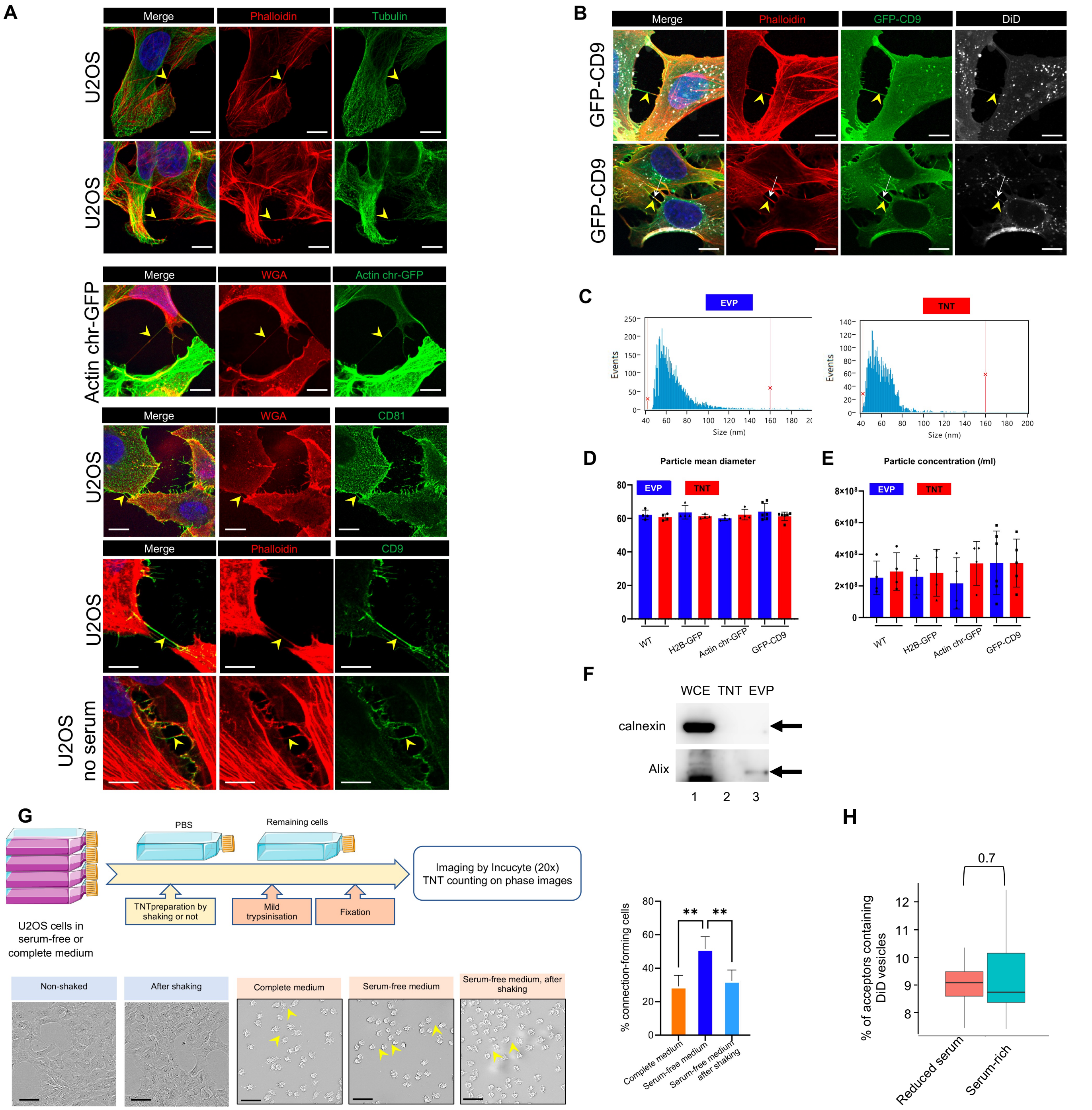

A
Merge
Phalloidin
Tubulin
U2OS
U2OS
Merge
WGA
Actin chr-GFP
Actin chr-GFP
Merge
WGA
CD81
U2OS
Merge
Phalloidin
CD9
U2OS
U2OS no serum
B
Merge
Phalloidin
GFP-CD9
DiD
GFP-CD9
GFP-CD9
C
EVP
TNT
Events
Size (nm)
D
Particle mean diameter
EVP
TNT
WT
H2B-GFP
Actin chr-GFP
GFP-CD9
E
Particle concentration (/ml)
EVP
TNT
WT
H2B-GFP
Actin chr-GFP
GFP-CD9
F
WCE TNT EVP
calnexin
Alix
1 2 3
G
PBS
Remaining cells
Imaging by Incucyte (20x) TNT counting on phase images
TNTpreparation by shaking or not
Mild trypsinisation
Fixation
U2OS cells in serum-free or complete medium
Non-shaked
After shaking
Complete medium
Serum-free medium
Serum-free medium, after shaking
% connection-forming cells
Complete medium
Serum-free medium
Serum-free medium after shaking
H
0.7
% of acceptors containing DiD vesicles
Reduced serum
Serum-rich

**Figure S1: Characterization of TNTs in U2OS cells**

A. Representative immunofluorescence of TNTs in U2OS cells, expressing actin (stained with phalloidin, red in the two first lanes) without or with tubulin (green, lanes 1 and 2). Actin-chromodoby-GFP is also expressed in TNTs (lane 3). Lanes 4 to 6 show representative images of CD81 and CD9 in TNTs of U2OS cells. In lane 6, cells were grown for 24h in the absence of serum before fixation. WGA (wheat germ agglutinin in red) labels cell surface of these non-permeabilized cells. The images in lanes 1 to 6 are respectively projections of 1, 3, 3, 1, 2 and 1 slices of each stack. The yellow arrowheads point to TNTs, connecting two cells and not attached to the substrate. The scale bars are 10 µm.

B. Two examples of TNTs in GFP-CD9 expressing U2OS cells. TNTs are identified as containing actin (phalloidin in red), and DiD labels intracellular vesicles, sometimes found inside TNT as in the second lane (white arrow). The yellow arrowheads point to TNTs, scale bars are 10 µm.

C. Representative examples of size distribution of EVPs and TNTs fractions.

D. Scatter dot plot representing mean size diameter of particles (with SD) of EVP and TNT fractions, depending on each cell line. At least 4 independent experiments were analyzed, each dot being one of them. No statistical difference was observed. Over 18 samples from all cell lines, mean particle diameter was 62.7 nm (SD : 3.8) for EVPs, 61.4 nm (SD 2.2) for TNTs.

E. Scatter dot plot representing mean particle concentration (with SD) of EVP and TNT fractions, depending on each cell line. At least 4 independent experiments were analyzed, each dot being one of them. No statistical difference was observed.

F. Representative western blots from 3 independent preparations of WCE (40 µg of proteins), TNT and EVP fractions (from about 20 $10^6$ cells), using the indicated antibodies successively.

G. Effect of shaking on U2OS (or actin chromobody-GFP expressing U2OS cells) cells and connections. The top diagram summarizes the flow of experiments aimed at visualizing and counting the connections between cells. Pictures were acquired using Incucyte (20x magnification) on flasks either with or witout shaking in serum-free medium (first two pictures), or after mild trypsinization of cells grown either in complete, or in serum-free medium, or remaining cells after shaking (pictures 3 to 5). Scale bars are 100 µm, arrowheads point to examples of cell-to-cell connections. The graph represents the percentage of connection-forming cells. Statistical analysis of 4 independent experiments was done by oneway Anova with Tukey post hoc correction (** p= 0.0052 and 0.0085 respectively)

H. U2OS cells are able to transfer vesicles when cultured in serum-rich or reduced serum-containing medium for 24 hours. Actin chromobody-GFP expressing U2OS cells were challenged with DiD for 30 min to label vesicles, next cocultured with mcherry-expressing U2OS as acceptors for 18 hours in complete or serum-free medium. Analysis was performed by FACS, boxplot shows percentage of acceptor cells positive for DiD (means respectively at 9.4 and 8.9%), statistical analysis is a pairwise comparison from 3 independent experiments (p=0.7).

### Figure S2

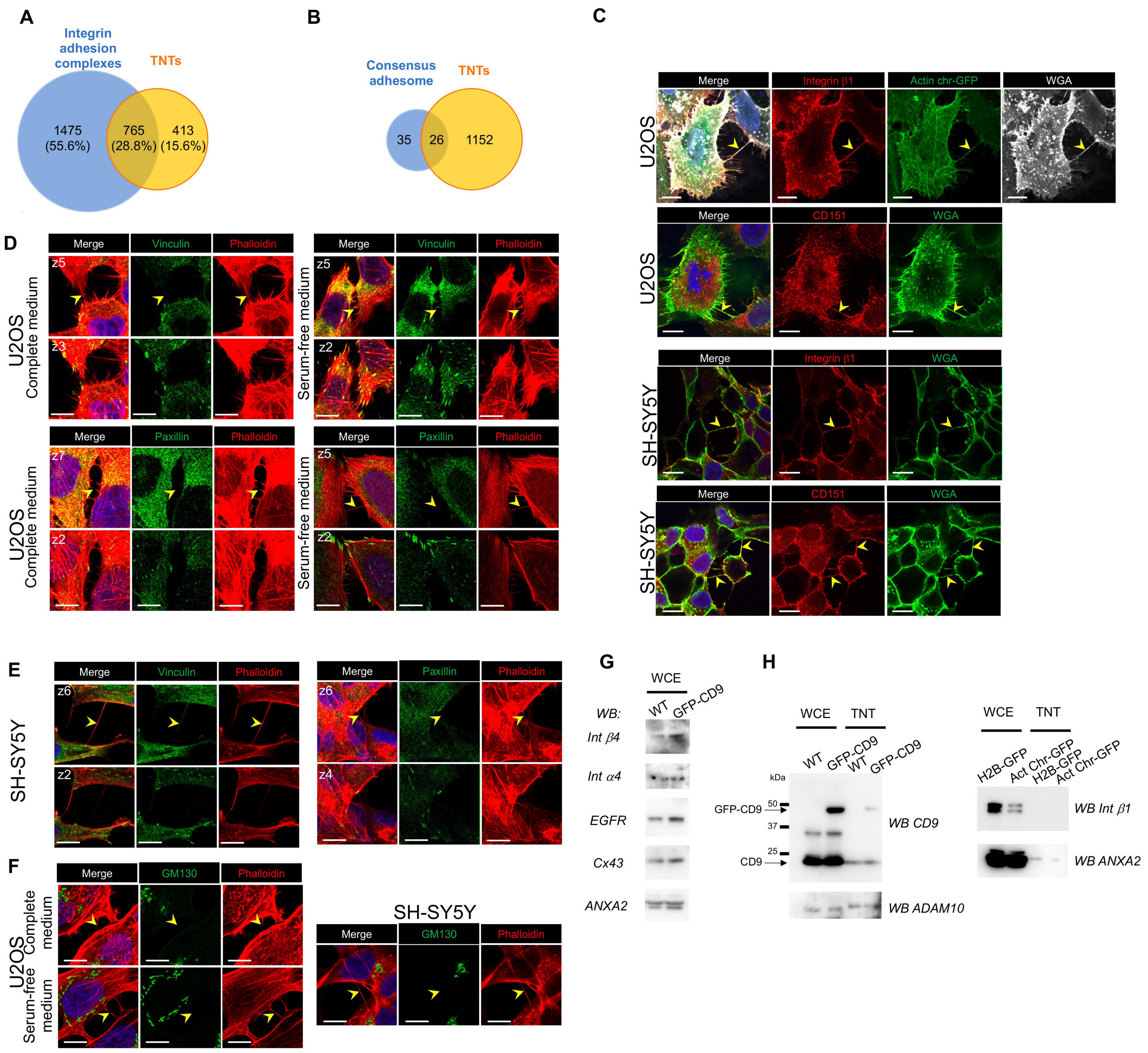
A
Integrin adhesion complexes
TNTs
1475 (55.6%)
765 (28.8%)
413 (15.6%)
B
Consensus adhesome
TNTs
35
26
1152
C
Merge
Integrin β1
Actin chr-GFP
WGA
U2OS
CD151
SH-SY5Y
D
Vinculin
Phalloidin
Paxillin
U2OS Complete medium
Serum-free medium
E
F
GM130
U2OS Complete medium
Serum-free medium
G
WCE
WB:
WT
GFP-CD9
Int β4
Int α4
EGFR
Cx43
ANXA2
H
WCE
TNT
kDa
GFP-CD9
CD9
50
37
25
WB CD9
WB ADAM10
H2B-GFP
Act Chr-GFP
WB Int β1
WB ANXA2

**Figure S2: Comparison of TNTome with Integrin adhesome and other cell proteins**

A. Venn diagram showing common and exclusive proteins between Integrin adhesion complexes (blue circle) and TNTs (yellow circle). The percentages refer to total proteins.
B. Venn diagram showing common and exclusive proteins between consensus adhesome (blue circle) and TNTs (yellow circle).
C. Representative immunofluorescence pictures showing expression of Integrin $\beta$1 and CD151 in TNTs of U2OS and SH-SY5Y as indicated on the left. Each picture is one upper slice of the stack, TNTs are further characterized by actin presence (actin chromobody-GFP, first lane) or WGA labeling. The yellow arrowheads point to TNTs, scale bars are 10 µm.
D. Representative immunofluorescence pictures showing labeling of Vinculin and Paxillin in U2OS cells cultured in complete medium or serum-free medium for 24h before fixation, as indicated on the left. For each labeling, bottom slice of the stack is shown on the bottom row (z number), upper slice shows a TNT, pointed with yellow arrowhead. Red is phalloidin staining, blue is DAPI in merge pictures; scale bars are 10 µm.
E. Representative immunofluorescence pictures showing labeling of Vinculin and Paxillin in SY-SY5Y cells, as in D.
F. Representative immunofluorescence pictures showing labeling of GM130 in U2OS cells (left, complete or serum-free medium), and SH-SY5Y cells. The yellow arrowheads point to TNTs, scale bars are 10 µm.
G. Expression of TM proteins in U2OS whole cell extracts (WCE), which are not in TNTome. WB from WT or GFP-CD9 expressing cells, incubated with the following antibodies as indicated on the left: Integrin $\beta$4 (Int $\beta$4), $\alpha$4 (Int $\alpha$4), EGFR and Connexin 43 (Cx43). Annexin A2 (ANXA2) is used as loading control. WCE from all cell lines have been tested three times, two WCE are shown.
H. Comparative expression of proteins in WCE and TNTs from various U2OS cell lines. Left, CD9, GFP-CD9 and ADAM10 are compared in WT and GFP-CD9 expressing U2OS cells (using non reducing gels). Right, Int $\beta$1 and ANXA2 are compared in H2B-GFP and Actin chromobodies-expressing cells (gels in reducing conditions).

### Figure S3

A

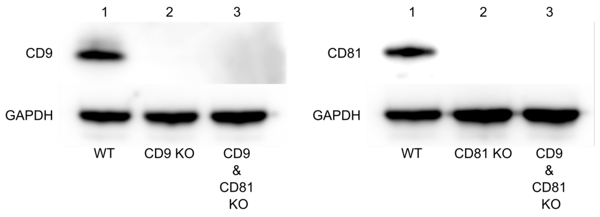


B

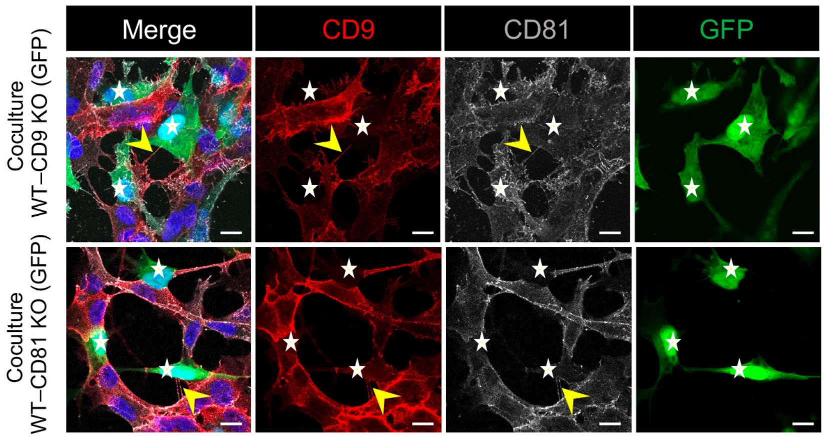


C

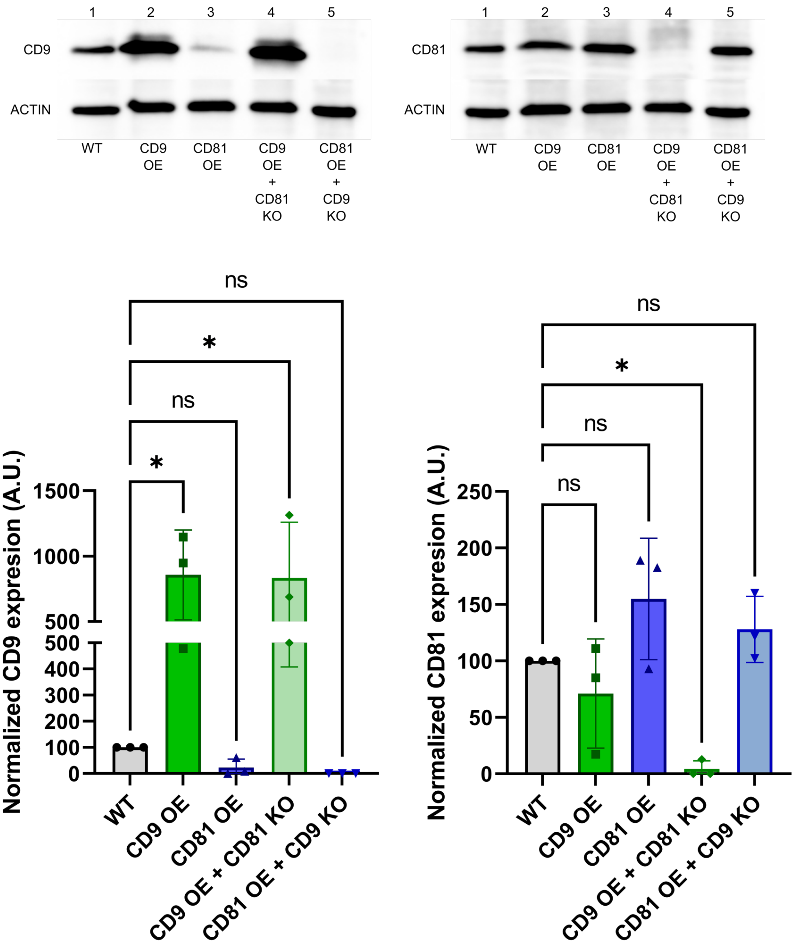

**Figure S3: Characterization of cells KO and OE for CD9 or CD81.**

A. Representative WB of the tetraspanin KO experiments blotted with CD9 (top left blot) or CD81 (top right blot) and GAPDH specific antibodies sequentially. Lines of the top left blot correspond to 1: WT cells; 2: CD9 KO cells; 3: CD9 & CD81 KO cells. Lines of the top right blot correspond to 1: WT cells; 2: CD81 KO cells; 3: CD9 & CD81 KO cells.

B. Representative immunofluorescence of coculture of WT SH-SY5Y cells with either CD9 KO cells (first row) or CD81 KO cells (second row), both expressing GFP. Anti-CD9 antibodies are in red (Alexa 546 secondary antibodies), anti-CD81 (Alexa 633 secondary antibodies) are in grey. KO cells (expressing GFP) are indicated by white stars, yellow arrowheads show TNTs, scale bars are 10 µm.

C. Quantification of the total amount of CD9 or CD81 from various cell extracts, blotted with CD9 (left blot) or CD81 (right blot) and actin specific antibodies sequentially. Lines of the left blot correspond to 1: WT cells; 2: CD9 OE cells; 3: CD81 OE cells; 4: CD9 OE + CD81 KO cells; CD81 OE + CD9 KO cells. Lines of the right blot correspond to 1: WT cells; 2: CD9 OE cells; 3: CD81 OE cells; 4: CD9 OE + CD81 KO cells; CD81 OE + CD9 KO cells. The graphs below show the relative expression of CD9 (bottom left) or CD81 (bottom right) in arbitrary units (A.U.). Bottom left: normalized expression of CD9 in WT cells (100%), CD9 OE cells (857.1% ± 343.5), CD81 OE cells (23.1% ± 32.7), CD9 OE + CD81 KO cells (833.4% ± 425.7) and CD81 OE + CD9 KO cells (0% ± 0) corresponding to the measurement of 3 independent WBs. Bottom right: normalized expression of CD81 in WT cells (100%), CD9 OE cells (71.1% ± 48.4), CD81 OE cells (154.9% ± 53.7), CD9 OE + CD81 KO cells (4.3% ± 7.4) and CD81 OE + CD9 KO cells (128% ± 29.3) corresponding to the measurement of 3 independent WBs. Statistical analysis was performed using Anova with Dunnett post-hoc test (ns: $p>0.05$, *: $p<0.05$).

### Figure S4

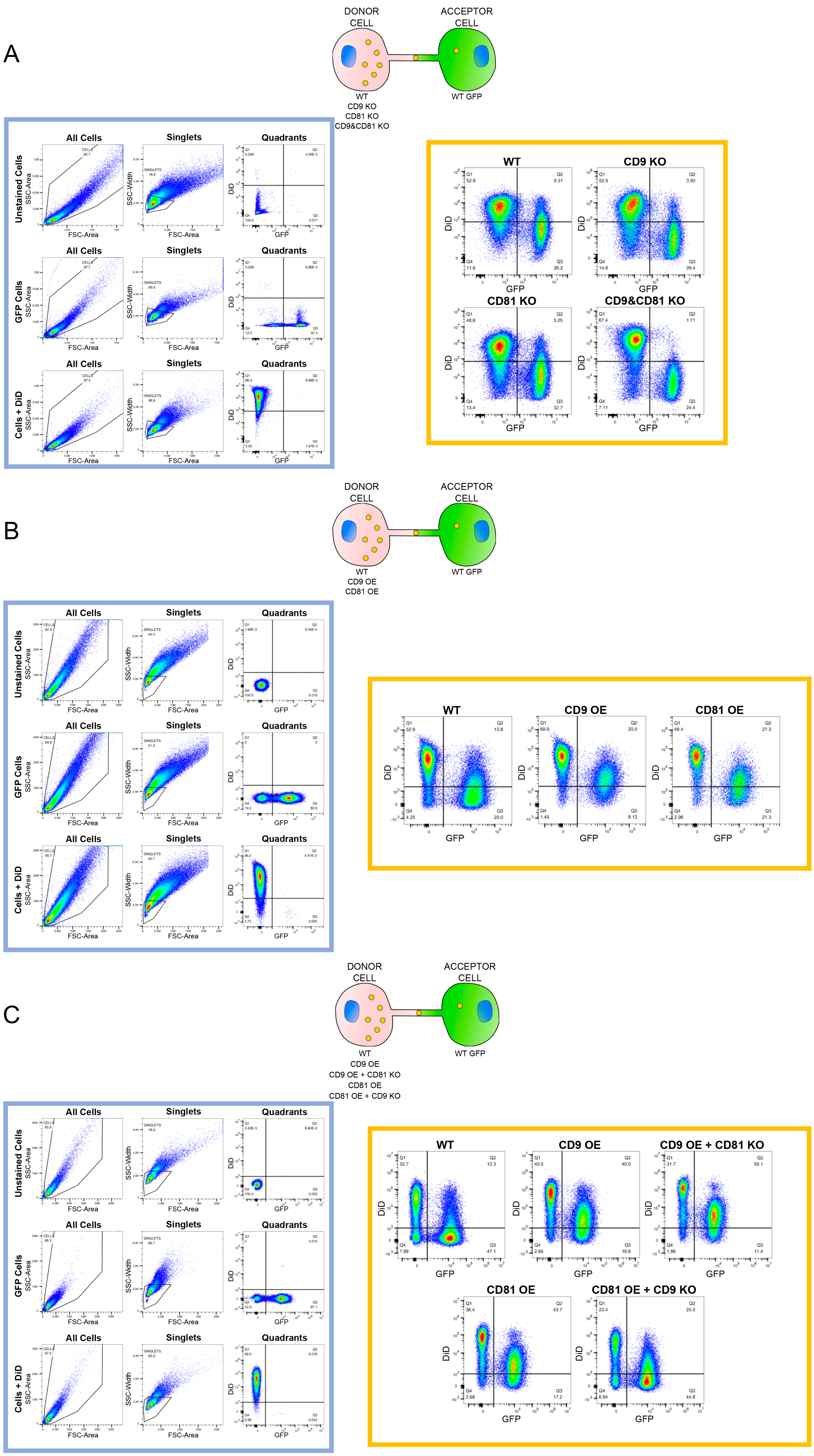
A
DONOR CELL
ACCEPTOR CELL
WT
CD9 KO
CD81 KO
CD9&CD81 KO
WT GFP
All Cells
Singlets
Quadrants
Unstained Cells
GFP Cells
Cells + DiD
SSC-Area
SSC-Width
FSC-Area
DiD
GFP
WT
CD9 KO
CD81 KO
CD9&CD81 KO
B
DONOR CELL
ACCEPTOR CELL
WT
CD9 OE
CD81 OE
WT GFP
WT
CD9 OE
CD81 OE
C
DONOR CELL
ACCEPTOR CELL
WT
CD9 OE
CD9 OE + CD81 KO
CD81 OE
CD81 OE + CD9 KO
WT GFP
WT
CD9 OE
CD9 OE + CD81 KO
CD81 OE
CD81 OE + CD9 KO

**Figure S4: Gate strategy and representative results of all the cocultures**

Schematics of the cocultures are on top of each panel. Gates strategies are framed in blue: WT unstained cells, GFP cells and DiD+ cells were gated to select cells and exclude cellular debris, plotting SSC-Area (Y-axis) with FSC-Area (X-axis) (All Cells). Within the All Cells gate we selected for single cells to exclude doublets, plotting SSC-Width (Y-axis) with FSC-Area (X-axis) (Singlets). Within the Singlets gate the quadrants for DiD (Y-axis) and GFP (X-axis) were placed according to the position of the negative or positive populations.
Framed in orange are representative plots of the signal of DiD-labeled vesicles (Y-axis) and GFP-labeled acceptor cells (X-axis). The second quadrant (Q2) represents the % of double positive cells and thus the acceptor cells that have received vesicles.

A. Coculture of tetraspanins KO cells used as donors (WT, CD9 KO, CD81 KO and CD9&CD81 KO) loaded with DiD to stain the vesicles with GFP WT cells
B. Coculture between tetraspanin OE cells ( WT, CD9 OE, CD81 OE) used as donors and WT GFP cells used as acceptors.
C. Coculture between tetraspanin OE + KO cells used as donors (WT, CD9 OE, CD9 OE + CD81 KO, CD81 OE and CD81 OE + CD9 KO) loaded with DiD to stain the vesicles and WT GFP cells used as acceptors.

### Figure S5

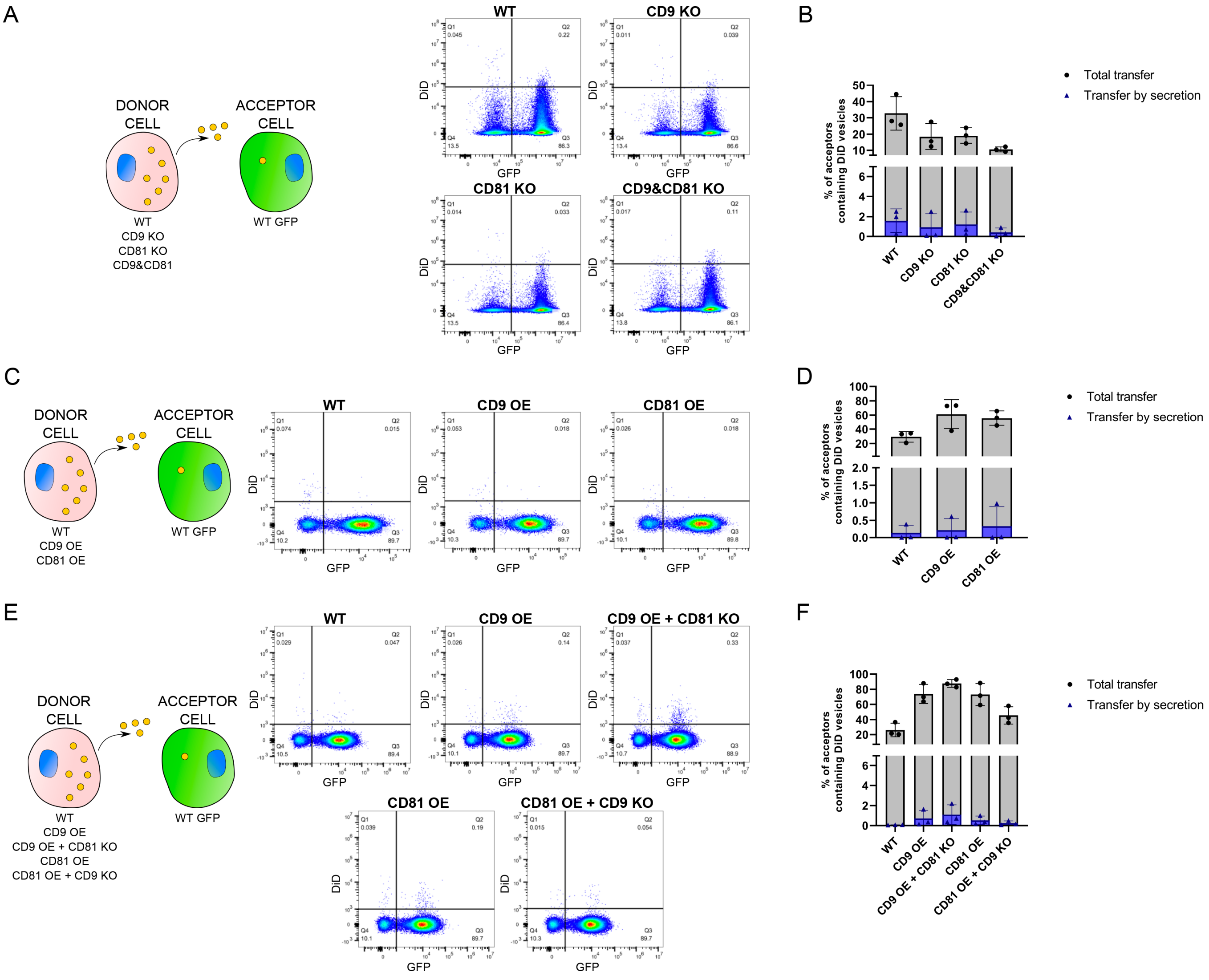

A
DONOR CELL
ACCEPTOR CELL
WT
CD9 KO
CD81 KO
CD9&CD81
WT GFP
WT
CD9 KO
CD81 KO
CD9&CD81 KO
DiD
GFP
B
Total transfer
Transfer by secretion
% of acceptors containing DiD vesicles
WT
CD9 KO
CD81 KO
CD9&CD81 KO
C
DONOR CELL
ACCEPTOR CELL
WT
CD9 OE
CD81 OE
WT GFP
WT
CD9 OE
CD81 OE
DiD
GFP
D
Total transfer
Transfer by secretion
% of acceptors containing DiD vesicles
WT
CD9 OE
CD81 OE
E
DONOR CELL
ACCEPTOR CELL
WT
CD9 OE
CD9 OE + CD81 KO
CD81 OE
CD81 OE + CD9 KO
WT GFP
WT
CD9 OE
CD9 OE + CD81 KO
CD81 OE
CD81 OE + CD9 KO
DiD
GFP
F
Total transfer
Transfer by secretion
% of acceptors containing DiD vesicles
WT
CD9 OE
CD9 OE + CD81 KO
CD81 OE
CD81 OE + CD9 KO

**Figure S5: Measurement of transfer by secretion in all the coculture experiments**

Schematics of all experiments are shown on the left.

A. Secretion control in the coculture between tetraspanin KO cells used as donors loaded with DiD to stain the vesicles and WT GFP cells used as acceptors. Representative plots between the signal of DiD-labeled vesicles (Y-axis) and GFP-labeled acceptor cells (X-axis) of vesicle transfer by secretion in cells: WT, CD9 KO, CD81 KO and CD9&CD81 KO. The second quadrant (Q2) represents the % of double positive cells and thus the acceptor cells that have received vesicles.

B. Graph showing the % of acceptor cells containing DiD vesicles from the coculture with tetraspanin KO cells used as donors, total and secretion-dependent. Total transfer is the same as in Figure 3F, the percentage of transfer by secretion is shown and always minor and not significantly different between conditions. For secretion, mean and standard deviation of the % of acceptor cells containing DiD vesicles are: WT = 1.6 ± 1.18; CD9 KO = 0.9 ± 1.38; CD81 KO = 1.2 ± 1.26; CD9 & CD81 KO = 0.4 ± 0.44 for N=3.

C. Secretion control in the coculture between tetraspanin OE cells used as donors loaded with DiD to stain the vesicles and WT GFP cells used as acceptors. Representative plots between the signal of DiD-labeled vesicles (Y-axis) and GFP-labeled acceptor cells (X-axis) of vesicle transfer by secretion in cells: WT, CD9 OE and CD81 OE. The second quadrant (Q2) represents the % of double positive cells and thus the acceptor cells that have received vesicles.

D. Graph corresponding to the % of acceptor cells containing DiD vesicles from the coculture with tetraspanin OE cells used as donors, total and secretion-dependent. Total transfer is the same as in Figure 3G, the percentage of transfer by secretion is shown and always minor and not significantly different between conditions. For secretion, mean and standard deviation of the % of acceptor cells containing DiD vesicles: WT = 0.14 ± 0.22; CD9 OE = 0.21 ± 0.34; CD81 OE = 0.33 ± 0.56 for N=3.

E. Secretion control in the coculture between tetraspanin OE + KO cells used as donors loaded with DiD to stain the vesicles and WT GFP cells used as acceptors. Representative plots between the signal of DiD-labeled vesicles (Y-axis) and GFP-labeled acceptor cells (X-axis) of vesicle transfer by secretion in cells: WT, CD9 OE, CD9 OE + CD81 KO, CD81 OE and CD81 OE + CD9 KO. The second quadrant (Q2) represents the % of double positive cells and thus the acceptor cells that have received vesicles.

F. Graph corresponding to the % of acceptor cells containing DiD vesicles from the coculture with tetraspanin OE + KO cells, total and secretion-dependent. Total transfer is the same as in Figure 5C, the percentage of transfer by secretion is shown and always minor and not significantly different between conditions. For secretion, mean and standard deviation of the % of acceptor cells containing DiD vesicles: WT = 0.07 ± 0.02; CD9 OE = 0.71 ± 0.80; CD9 OE + CD81 KO = 1.09 ± 0.97; CD81 OE = 0.52 ± 0.42; CD81 OE + CD9 KO = 0.24 ± 0.22 for N=3.

### Figure S6

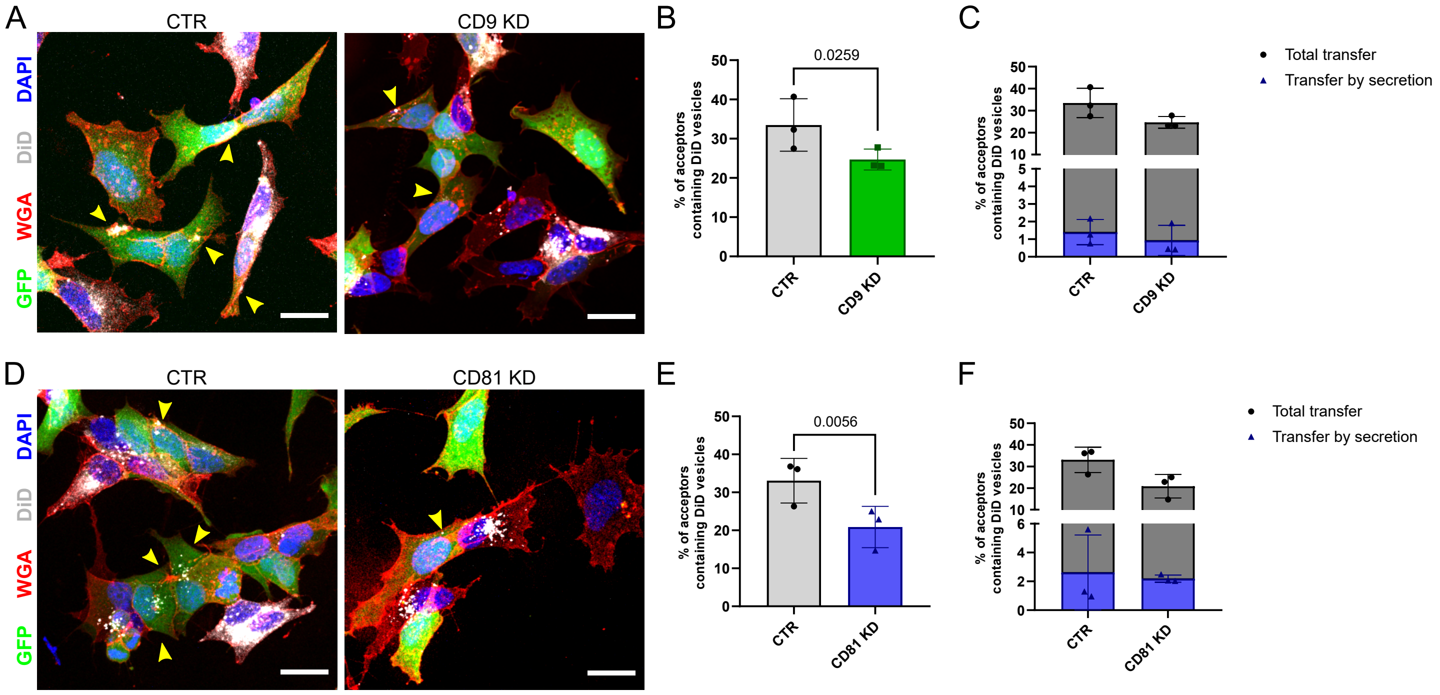

A
CTR
CD9 KD
DAPI
DiD
WGA
GFP
B
0.0259
% of acceptors containing DiD vesicles
CTR
CD9 KD
C
Total transfer
Transfer by secretion
% of acceptors containing DiD vesicles
CTR
CD9 KD
D
CTR
CD81 KD
DAPI
DiD
WGA
GFP
E
0.0056
% of acceptors containing DiD vesicles
CTR
CD81 KD
F
Total transfer
Transfer by secretion
% of acceptors containing DiD vesicles
CTR
CD81 KD

**Figure S6: Coculture of DiD-treated WT, CD9KD or CD81KD donor cells with GFP-expressing cells as acceptor, and analysis of transfer by microscopy.**

A. Representative confocal micrograph of the cocultures between KD control cells (CTR) or the KD of CD9 (CD9 KD). Human siRNA for CD9 (sequence: GAGCATCTTCGAGCAAGAA) or non-targeting siRNA (CTR from Origene) were transiently transfected into cells using Lipofectamine RNAimax (Invitrogen), in accordance with the manufacturer's instructions. Experiments were performed 48 hours following transfection. CTR or CD9 KD cells were used as donors and therefore loaded with DiD (white) to stain the vesicles and they were mixed in a ratio 1:1 with acceptors cells that were expressing GFP (green). 24 hours later, cells were fixed and stained with WGA (red) to stain the membranes and DAPI (blue) to stain the nuclei. Yellow arrowheads mark acceptor cells containing DiD-vesicles. The scale bars correspond to 10 µm.
B. Graphs of the coculture represented in A, showing the % of acceptor cells containing DiD vesicles corresponding to the total transfer from 3 independent experiments. Mean ± SD are: CTR = 33.5 ± 6.68; CD9 KD = 24.7 ± 2.68.
C. Same graph as in B corresponding to the % of acceptor cells containing DiD vesicles of the coculture represented in A, including the part due to transfer by secretion. Secretion-dependent transfer is minor and not significantly different between the conditions, mean ± SD are: CTR = 1.4 ± 0.72; CD9 KD = 0.92 ± 0.86 for N=3.
D. Representative confocal micrograph of the cocultures between KD control cells (CTR) or the KD of CD81 (CD81 KD). siRNA sequence targeting human CD81 is: GCACCAAGUGCAUCAAGUA. The experiment was perform as described in A. CTR or CD81 KD cells were used as donors and therefore loaded with DiD (white) to stain the vesicles and they were mixed in a ratio 1:1 with acceptors cells that were expressing GFP (green). Yellow arrowheads mark acceptor cells containing DiD-vesicles. The scale bars correspond to 10 µm.
E. Graphs of the coculture represented in D, showing the % of acceptor cells containing DiD vesicles corresponding to the total transfer from 3 independent experiments. Mean ± SD are: CTR = 33.1 ± 5.87; CD81 KD = 20.9 ± 5.44.
F. Same graph as in E corresponding to the % of acceptor cells containing DiD vesicles, including the part due to transfer by secretion of the coculture represented in D. Secretion-dependent transfer is minor and not significantly different between the conditions, mean ± SD are: CTR = 2.6 ± 2.6; CD81 KD = 2.2 ± 0.25 for N=3.

### Figure S7

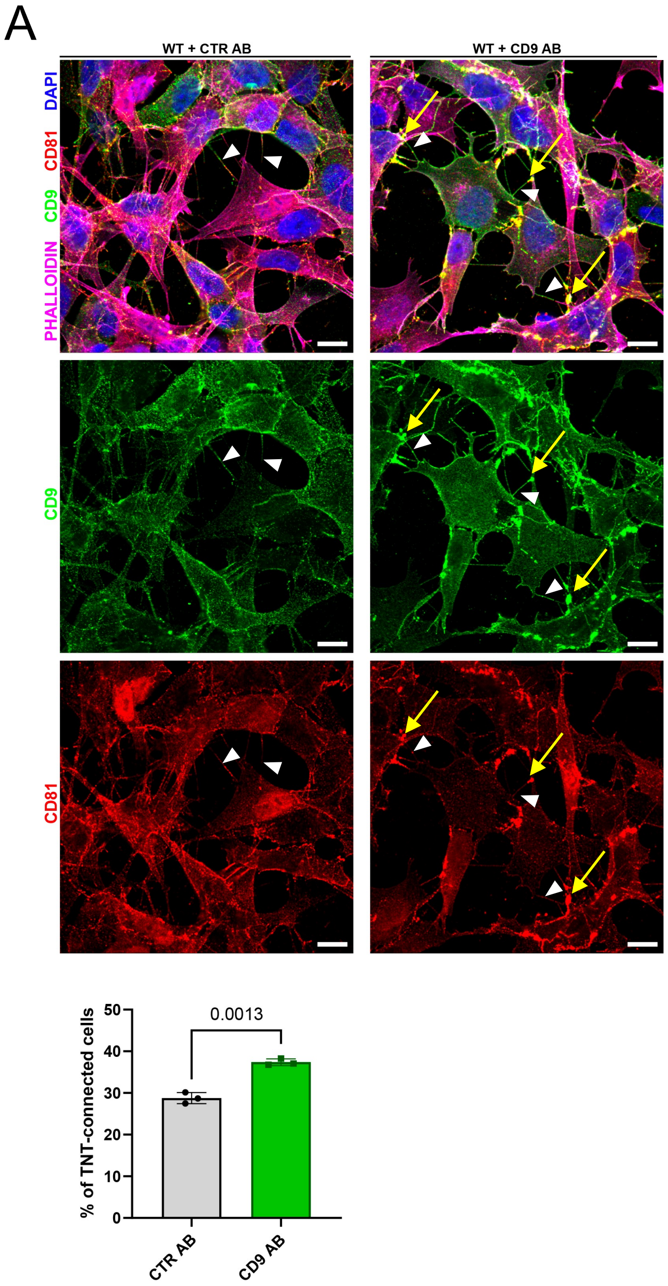
A
WT + CTR AB
WT + CD9 AB
DAPI
CD81
CD9
PHALLOIDIN
CD9
CD81
0.0013
% of TNT-connected cells
50
40
30
20
10
0
CTR AB
CD9 AB

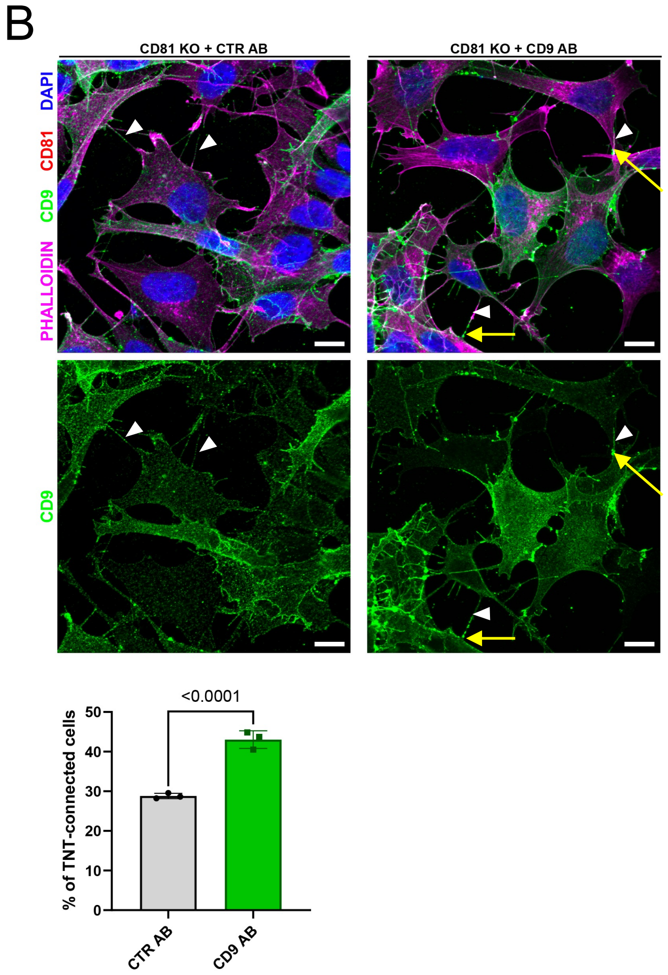
B
CD81 KO + CTR AB
CD81 KO + CD9 AB
DAPI
CD81
CD9
PHALLOIDIN
CD9
<0.0001
% of TNT-connected cells
50
40
30
20
10
0
CTR AB
CD9 AB

**Figure S7: CD9 vs. CTR antibody treatment in WT and CD81 KO cells**

A. A. Immunofluorescence of CD9 (green) and CD81 (red) in SH-SY5Y WT cells treated with control antibodies (CTR AB) or with antibodies anti-CD9 (CD9 AB) for 2 hours followed by PFA fixation and incubation with anti-CD81 (for CD9 AB samples) or anti-CD81 + anti-CD9 (for CTR AB samples) and appropriate fluorescent secondary antibodies. This representative image corresponds to the 3rd to 4th slices of a stack comprising 12 slices for CTR AB and to the 3rd to 4th slices of a stack comprising 11 slices for CD9 AB. Cells were also stained with phalloidin (magenta) and DAPI (blue) to visualize actin and nuclei respectively. White arrowheads show TNTs, yellow arrows highlight CD9 and CD81 clustering area. The scale bars correspond to 10 µm. Below is the graph of the % of TNT-connected cells. Mean ± SD (N=3) are: CTR AB = 28.8 ± 1.32; CD9 AB = 37.4 ± 0.80.

B. Immunofluorescence of CD9 (green) in SH-SY5Y CD81 KO cells treated with control antibodies (CTR AB) or with antibodies anti-CD9 (CD9 AB) for 2 hours followed by PFA fixation and incubation with anti-CD9 (for CTR AB samples) and appropriate fluorescent secondary antibodies. This representative image corresponds to the 3rd to 4th slices of a stack comprising 10 slices for CTR AB and to the 5th to 6th slices of a stack comprising 12 slices for CD9 AB. Cells were also stained with phalloidin (magenta) and DAPI (blue) to visualize actin and nuclei respectively. White arrowheads show TNTs, yellow arrows highlight CD9 clustering area Scale bars are 10 µm. Below is the graph of the % of TNT-connected cells. Mean ± SD (N=3): CTR AB = 28.8 ± 0.68; CD9 AB = 43 ± 2.23.
